# Accelerated Evolution of Context-Dependent Gene Expression in Primate Immune Cells

**DOI:** 10.64898/2026.09.23.753790

**Authors:** Arun Durvasula, Amy Longtin, Audrey M. Arner, Kathrin Köhler, Iker Rivas-González, Jenny Tung, Genevieve Housman, Amanda J. Lea

**Author notes:** Correspondence should be addressed to A.D., G.H., or A.J.L. These authors contributed equally (Genevieve Housman, Amanda J. Lea).

## Abstract

Regulatory variation is widely recognized as a driver of evolution within and between species, including in the primate lineage. Evidence for potentially adaptive regulatory evolution in primates comes from both sequence-based tests for selection and cross-species comparisons of gene expression. However, comparative studies of gene expression have tended to focus on baseline cellular states, despite growing evidence that context-dependent gene regulation is important. Here, we investigate the evolution of gene expression in lymphoblastoid cell lines (LCLs) from 6 primate lineages (humans, chimpanzees, bonobos, western gorillas, Sumatran orangutans, and rhesus macaques) using data from both control and 5 perturbed cellular conditions (n=22-25 individuals per condition). Analyzing baseline expression data, we find that genes involved in immunity often exhibit lineage-specific transcriptional shifts, and that these genes occur near regions with lineage-specific sequence acceleration and lineage-specific epigenomic states. However, the subset of genes that respond to *in vitro* perturbations have evolved at significantly higher rates than environmentally-insensitive genes, a finding we replicate in an independent four-species dataset of primary white blood cells (n=26). Together, our findings support the argument that focusing on baseline states alone provides a biased view of gene regulatory evolution, and suggest that studying gene regulation in dynamic contexts can enrich our understanding of primate evolution.

## Introduction

Changes in gene regulation have long been proposed to play an important role in trait evolution within and between species [1–4]. In support of this hypothesis, a large body of work has identified positively selected regulatory mutations that shape morphology, physiology, and behavior across vertebrates [3,5–8]. For example, marine sticklebacks that have colonized freshwater lakes have evolved reduced skeletal armor relative to their freshwater counterparts through regulatory changes near *PITX1*, a developmental transcription factor [9–12]. Similarly, selection on regulatory variation controlling mesenchyme expression of the genes *ALX1* and *HMGA2* has shaped the evolution of beak morphology during the adaptive radiation of Darwin’s finches [13,14]. In primates specifically, many efforts have been made to link regulatory mutations to interspecific trait variation (e.g., [15–19], building on early arguments for the importance of gene regulation in comparisons between humans and chimpanzees [2]. However, compared to more tractable experimental systems [20,21], we still know relatively little about how evolution has shaped gene regulatory variation in the primate lineage [3,22].

To evaluate the evolution of primate gene regulation, researchers have typically relied on two approaches: sequence-based tests for selection and comparative analyses of gene expression. Sequence-based approaches have identified signatures of natural selection in regulatory regions within and between species (e.g., [23–26]), sometimes complemented with functional follow-up. For example, a recent analysis of genomes from 49 primate species identified a region of accelerated evolution in gibbons; this mutation modifies gene expression in the limbs and shoulders of transgenic mice and is a candidate contributor to this taxon’s strikingly elongated arms, which facilitate arboreal locomotion [27]. In parallel, comparative transcriptomic analyses have shown that gene expression divergence generally mirrors phylogeny [28–30]. However, for some genes and tissues (e.g., mammalian testes: [29–31]), the phylogenetic pattern is violated, potentially pointing to targets of selection. Some comparative studies have also fit evolutionary models to transcriptomic data to infer selection [17,29,32–37], suggesting that at least some gene expression levels evolve under selection (although technical concerns, such as species differences in tissue composition, can influence the reliability of these inferences [38]).

In combination, both sequence-based studies and comparative transcriptomic studies suggest that primate gene expression levels are generally constrained by purifying and stabilizing selection, but that positive selection can act to shape gene expression in a gene-, pathway-, or tissue-specific manner [31,39–42]. For example, previous work has shown that genes expressed broadly across tissues tend to evolve more slowly, while genes with more narrow tissue-specific functions evolve faster. This pattern is consistent with the idea that selection is more effective when targeting tissue-specific regulatory regions because it avoids costly pleiotropy [30,40]. Notably, although the same argument applies to other forms of context-specificity, such as environment- or developmental timing-specific patterns of gene regulation, much less work has focused on these forms of context-specificity [19,22,43] than on cross-tissue comparisons [17,28,29,31,40].

This leaves an important gap, because growing evidence points toward environmentally-dependent gene regulation—where expression programs are activated by certain physiological challenges, signals, or perturbations—as an overlooked component of trait genetic architecture. For example, work in humans has emphasized that, while an estimated 80-90% of all disease-associated genetic variation (identified via genome-wide association studies) is non-coding [44], only ∼50% of these variants can be linked to gene expression variation in any well-studied tissue [45]. While there are multiple potential explanations for the incomplete overlap [46], one possibility is that disease-relevant variants are not active in baseline cell states [47]. Indeed, studies that have explicitly mapped genetic effects on gene expression in cells challenged with pathogens, drugs, hormones, and other stimuli have revealed the widespread existence of “environmentally-dependent” expression quantitative trait loci (eQTL) [48–52]. This sub-type of context-dependent eQTL exhibits greater overlap with GWAS hits for complex traits and diseases than eQTL mapped under baseline conditions [48,50,53]. Additionally, environmentally-dependent eQTL identified using immune-related perturbations are more strongly enriched for signatures of positive selection than eQTL that are present independent of perturbations [49,50,54,55], consistent with the idea that context-specific variants should be more evolvable due to their reduced potential for pleiotropic, deleterious effects [3,47,56,57]. This work argues that perturbed cell states are important for understanding regulatory evolution, motivating comparative studies that incorporate environmental challenges.

To address this gap, we profiled genome-wide gene expression levels in lymphoblastoid cell lines (LCLs) derived from human, chimpanzee, bonobo, western gorilla, Sumatran orangutan, and rhesus macaque. We focused on LCLs given their accessibility across species, the extensive evidence for positive selection on both innate and adaptive immune processes [19,43,58,59], and the long history of LCLs as models for understanding the genetics and evolution of gene expression in primates [60–62] (Supplemental Text). Additionally, LCLs represent a single cell type, allowing us to eliminate cell type composition as a confounding factor in our analysis [38]. We used an *in vitro* perturbation design, in which gene expression was measured in parallel in both baseline, unperturbed cells and following experimental challenges with diverse stimuli that include hormonal (aldosterone), nutrient (glucose), cell stressor (tunicamycin), and medically relevant chemicals (caffeine, acetaminophen). This design allows us to understand the broad impact of environmental sensitivity on gene expression evolution, rather than focusing *a priori* on individual responses, such as the immune response, which are already relatively well-studied.

Using the resulting dataset (n =169 transcriptomes from 25 individuals and 6 species), we applied a Brownian motion model to infer rates of gene expression evolution across the phylogeny and to identify lineages and genes where evolutionary changes have occurred [36]. We then compared evolutionary rates for genes that change gene expression after perturbation (environmentally-dependent genes”) versus those that do not (“environmentally-insensitive genes”). We predicted that environmentally-dependent genes evolve faster than environmentally-insensitive genes, both to keep up with changing environmental pressures (e.g., Red Queen dynamics [63]) and because context-specificity can minimize pleiotropy. Finally, we performed a replication study of our main findings using transcriptome data from baseline and stimulated primary immune cells collected from 26 humans, chimpanzees, baboons, and rhesus macaques by Hawash and colleagues [19]. Overall, our analyses suggest that environmentally-dependent genes represent an evolutionary distinct class with faster evolutionary rates than environmentally-insensitive genes. We suggest these differences are driven by a combination of relaxed constraint and lineage-specific adaptive evolution.

## Results

### Study and dataset overview

We measured gene expression in baseline (unperturbed) and environmentally perturbed lymphoblastoid cell lines (LCLs) from humans: *Homo sapiens* (n=9; final sample size varied from 5-9 across conditions), chimpanzees: *Pan troglodytes* (n=5), bonobos: *Pan paniscus* (n=2), western gorillas: *Gorilla gorilla* (n=2), Sumatran orangutans: *Pongo abelii* (n=2), and rhesus macaques: *Macaca mulatta* (n=5) (**Figure 1A**, **Table S1**). We exposed cells from each individual for 4 hours to two baseline control conditions (water and ethanol) and five perturbation conditions (aldosterone, caffeine, glucose, tunicamycin, and acetaminophen) (**Figure 1B**, **Table S2**). Following mRNA-seq data generation, quality control, and low-level processing (with special attention to considerations for cross-species comparison; see Methods and **Table S3**), we compiled expression values for 12,455 protein-coding genes across all species and conditions.

**Figure 1.**
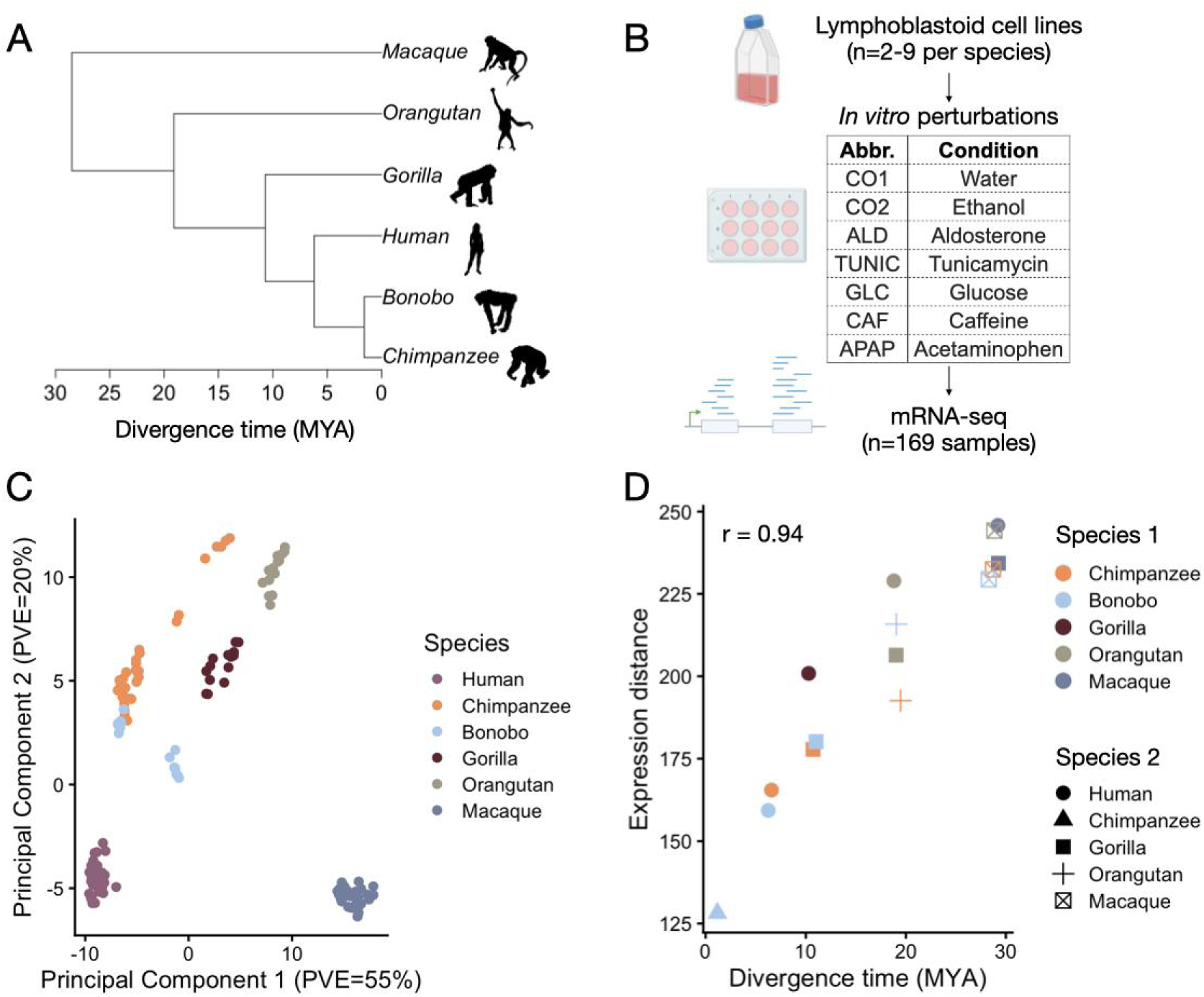
Study overview and gene expression data. A) Phylogeny of the species included in this study and B) overview of experimental design. LCLs from multiple species were exposed to control and treatment conditions to measure gene expression levels at baseline and in response to five different perturbations. Water was included as the vehicle control for glucose and caffeine. Ethanol was included as the vehicle control for acetaminophen, aldosterone, and tunicamycin. Evolutionary divergence times (million years ago, MYA) were obtained from [64,65] (see **Table S4**). Silhouette images were adapted from http://phylopic.org/, courtesy of T. Michael Keesey, Tony Hisgett, and Gareth Monger (http://creativecommons.org/licenses/by/3.0/). Additional images were obtained from BioRender. C) PCA of LCL gene expression data across species and treatments. See Figure S1 for PCA colored by treatments. PVE = percent variance explained. D) Comparison between divergence times (using estimates from [64,65] as in Panel A) and mean gene expression distance for each species pair, with gene expression distance calculated from a matrix of mean, voom-normalized expression values for each gene and species. Points are slightly jittered in the x-axis plane for visualization. The r value represents the correlation coefficient from a Mantel test. For all plots, we define taxonomic groupings using common names: human (*Homo sapiens*), chimpanzee (*Pan troglodytes*), bonobo (*Pan paniscus*), gorilla (*Gorilla gorilla*), orangutan (*Pongo abelii*), and macaque (*Macaca mulatta*).

Using these data, we first investigated whether gene expression levels globally recapitulated the known primate phylogeny, as expected based on prior work [28–30]. To do so, we performed principal component analysis (PCA) on all 169 transcriptomes (**Figure 1C**). The first principal component explained 56% of the overall variance in the data and separated all apes from the rhesus macaques (**Figure S1**), with subsequent PCs capturing variation within apes. As expected, transcriptomic variation more strongly mirrored the evolutionary history of the species rather than differences between conditions (**Figure S1**). Hierarchical clustering of the mean gene expression levels per species recapitulated the expected phylogeny (**Figure S1**) and pairwise estimates of genetic distance between species (from [64]) were significantly correlated with transcriptomic distances between species (Mantel test r =0.945, p=2.78×10^−3^) (**Figure 1D**).

### Heterogenous rates of gene expression evolution across great apes

We next modeled the evolution of gene expression levels across the phylogeny using the Computational Analysis of Gene Expression Evolution (CAGEE) approach [36,37]. In brief, CAGEE uses transcriptomic data for each taxon to estimate genome-wide evolutionary rates, which can be interpreted as the variance of the Brownian motion process per million years across log-transformed gene expression values. In addition, CAGEE identifies genes with credible increases or decreases in expression along a focal branch relative to the ancestor and infers differences in evolutionary rates between *a priori* defined gene lists.

Before turning our attention to the empirical data, we first performed simulations, conditional on the known phylogeny (**Table S4**), to understand power (**Figure S2**). Our first set of simulations focused on 7 scenarios: a null scenario in which a single rate is shared across the tree and six scenarios in which the evolutionary rate changes in one lineage, rotating through human, chimpanzee, bonobo, gorilla, orangutan, and macaque. Our second set of simulations focused on five scenarios: (i) a null scenario in which a single rate is shared across the tree, and four alternative scenarios in which the expression rate changes in a particular clade. These clades were defined as (ii) the chimpanzee-bonobo lineage (i.e., the genus *Pan*), (iii) the human-chimpanzee-bonobo (Hominini) lineage, (iv) the human-chimpanzee-bonobo-gorilla (Homininae) lineage, or (v) the human-chimpanzee-bonobo-gorilla-orangutan (Hominidae or great ape) lineage.

Our simulations showed that we had substantial power to detect species-specific rate changes in relatively short branches (chimpanzee, bonobo, and human), but weaker power to detect lineage-specific rate increases on longer branches (orangutan and macaque) (**Figure 2A, Table S5**). For example, we estimate 99% power to detect a 1.5x increase in evolutionary rate on the human lineage and 0% power to detect the same increase in orangutan and macaque. We found that we also had substantial power to detect rate lineage-specific increases that affected great ape clades (**Figure 2B**). For example, we estimate 100% power to detect a 1.5x increase in evolutionary rates on the human-chimpanzee-bonobo lineage, as well as the human-chimpanzee-bonobo-gorilla lineage. We generally find lower power to detect decreases than increases in the evolutionary rate for chimpanzees and bonobos, but greater power for the other lineages (**Figure 2**, **Table S5**).

**Figure 2.**
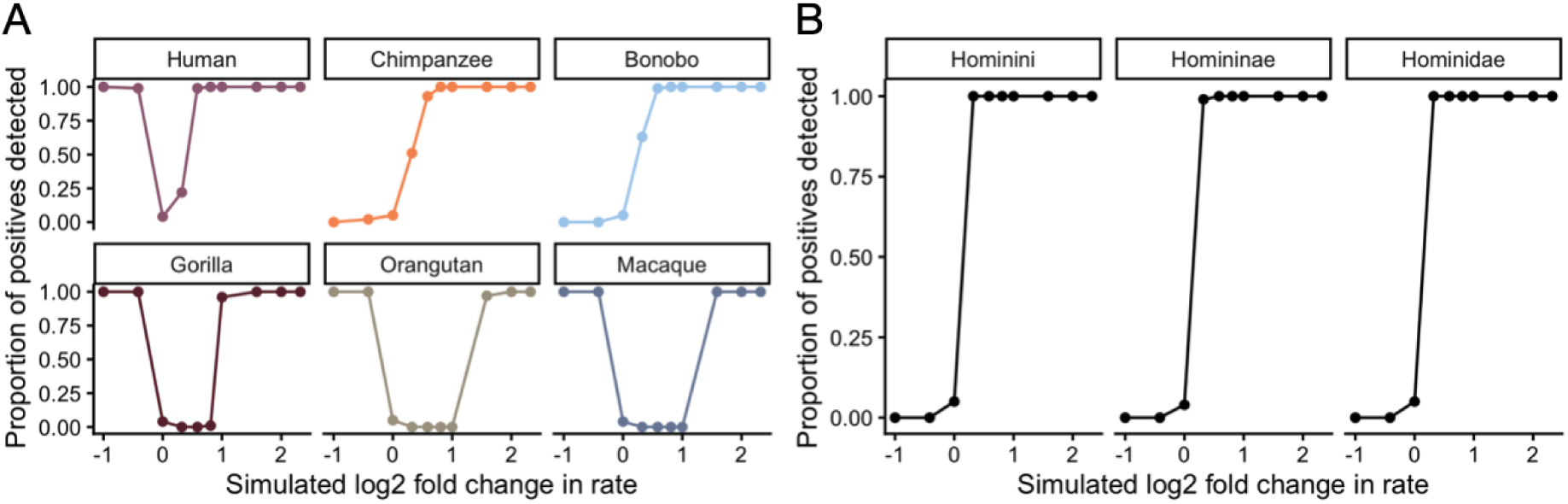
Power to detect accelerated gene expression evolution. Proportion of positives detected in 100 replicate simulations of accelerated evolutionary rates. For each scenario, we simulated expression rate increases from the base value of 0.0001 by the factor specified on the x-axis; rate increases were simulated in the lineages noted in the panel legends, which focused on A) terminal branches or B) clades. Terminal branch taxonomic groupings using common names: human (*Homo sapiens*), chimpanzee (*Pan troglodytes*), bonobo (*Pan paniscus*), gorilla (*Gorilla gorilla*), orangutan (*Pongo abelii*), and macaque (*Macaca mulatta*). Clades include Hominini (human-chimpanzee-bonobo lineage), Homininae (human-chimpanzee-bonobo-gorilla lineage), and Hominidae (human-chimpanzee-bonobo-gorilla-orangutan lineage).

In addition to estimating genome-wide rates of gene expression evolution, CAGEE also produces gene-specific estimates of expression level changes across branches of the phylogeny. In null simulations with no evolutionary rate change and inference under the single rate model, we observed a significant negative correlation between the estimated change in expression levels between sister taxa. For example, the correlation of gene-specific estimates for the chimpanzee and bonobo lineages is −0.11 (Spearman’s correlation, p<2×10^16^), and the correlation of gene-specific estimates for the human and the ancestral *Pan* lineage (the lineage from the common ancestor between humans, chimpanzees, and bonobos to the common ancestor of chimpanzees and bonobos) is −0.32 (p<2×10^16^). This pattern likely reflects the fact that changes in gene expression are relative to other taxa; an increase in gene expression in one lineage is relative to a decrease in another lineage. Therefore, comparisons of strictly increasing or decreasing genes between sister taxa should not be made. In addition, we found that the number of accelerated genes detected across taxa was substantially different even if the same overall rate increase was simulated. For example, with a 1.75x rate increase simulated on a single lineage, we found 1,632 accelerated genes in bonobo, 1,572 accelerated genes in chimpanzee, and 722 accelerated genes in macaque. These results are concordant with the recommendation of Bertram et al that comparisons of the total number of significantly accelerated or decelerated genes across taxa with different branch lengths should not be made [36].

Turning to the empirical data, we next investigated whether gene expression evolution varies in tempo across species using data from the baseline conditions (water and ethanol). To do so, we compared 12 different models of gene expression evolution: a single-rate shared across all lineages, and 10 two-rate models (6 species-specific and 4 clade-specific), as well as a model where all branches, internal and external, were allowed their own rates. We found strong support for two-rate models, where a different evolutionary rate characterizes a single species or clade relative to the rest of the phylogeny. Consistent with power differences revealed by our simulations, models with accelerated gene expression evolution for the chimpanzee lineage, bonobo lineage, and combined *Pan* lineage had the highest likelihoods relative to the single rate model. For example, the difference in log-likelihood between a model where the chimpanzee lineage had an accelerated rate compared to the single lineage model was 5,237 (**Table S6**). The second and third best fitting models were 1) a shared rate between chimpanzee and bonobo (Δ log-likelihood compared to single rate model: 4,470) and 2) a bonobo-specific rate (Δ log-likelihood compared to single rate model: 4,256). All three top models supported substantial acceleration in *Pan*: either a ∼7-fold higher rate of transcriptomic evolution for the chimpanzee lineage compared to the rest of the phylogeny (0.169 for chimpanzee, 0.024 for the rest of the tree; **Table S6**), a ∼6-fold acceleration for bonobo relative to the rest of the tree (0.15 versus 0.025), or a ∼3.4-fold acceleration when comparing *Pan* to all other species (0.074 versus 0.022). We observed very similar results when using gene expression values in the baseline ethanol condition. The top three models were 1) chimpanzee-specific rate (Δ log-likelihood compared to single rate model: 5,017; 6x increase in rate), 2) bonobo-specific rate (Δlog-likelihood compared to single rate model: 4,339; 3x increase in rate), and 3) Pan-specific rate (Δ log-likelihood compared to single rate model: 4,020; 3x increase in rate) (**Table S6**).

### Accelerated gene expression evolution across the ape phylogeny tracks immune function and sequence evolution

While our power to detect genes with credible increases in gene expression levels is uneven across the tree, our issues stem from high false negative rather than false positive rates for certain species, motivating us to investigate the genes we did detect. Specifically, for each of our two-rate species-specific models, in which the rate was allowed to vary along a terminal branch relative to the rest of the tree, we extracted genes for which CAGEE identified a credible expression difference on that branch relative to the immediate parent node (e.g., human compared to the *Homo*-*Pan* ancestor, chimpanzee compared to the *Pan* ancestor, etc.). Because the tree structure creates variation in power to detect credible gene expression differences between terminal and parent nodes (**Figure 2**), we identify different numbers of genes across species (range of genes with credible increases=0-1652, range of genes with credible decreases=421-2166; **Figure S3**, **Table S7**). Many of these genes overlapped across taxa (**Figure S4**), especially between closely related species (e.g., 90% of genes with credible increases or decreases in expression in chimpanzees were also shared with bonobos; Fisher’s exact test: odds=137.7, p<10^−16^). To check these results against an independent method, we compared the significant human-accelerated and chimpanzee-accelerated genes identified in our CAGEE analysis to those previously identified by Khan and colleagues from human, chimpanzee, and rhesus macaque LCL transcriptome data [42] We also identify a significant overlap between data sets for both human- and chimpanzee-accelerated genes in this comparison (Fisher’s exact test: odds=2.54 and 1.86, p-value = 1.30×10^−6^ and 4.26×10^−3^, respectively).

To obtain biological insight into the genes that exhibit rate changes, we asked whether the genes with credible increases in expression levels in a given species were enriched within particular biological pathways or processes relative to the background set of all genes expressed in LCLs (see **Figure S5** for parallel analyses that include genes with credible increases or decreases as the query set). Across taxa with non-zero numbers of accelerated genes, we found shared enrichment for immune-related processes, although not individual immune-related pathways. For example, human-accelerated genes were enriched for immune-response activating cell surface receptor signaling pathway (adjusted p-value=0.045) and leukocyte activation (adjusted p-value=0.045). Chimpanzee-accelerated genes were enriched for MHC protein complex assembly (adjusted p-value=1.02×10^−7^) and antigen processing and presentation (adjusted p-value=5.19×10^−4^) (**Figure 3A**, **Table S8**). Bonobo-accelerated genes were enriched for functions such as mononuclear cell differentiation (adjusted p-value=1.39×10^−4^) and B cell activation (adjusted p-value=1.44×10^−4^), and gorilla-accelerated genes were enriched for leukocyte migration (adjusted p-value=1.35×10^−5^) and lymphocyte chemotaxis (adjusted p-value=1.88×10^−5^). No significantly enriched gene ontology categories were identified for orangutan-accelerated genes.

**Figure 3.**
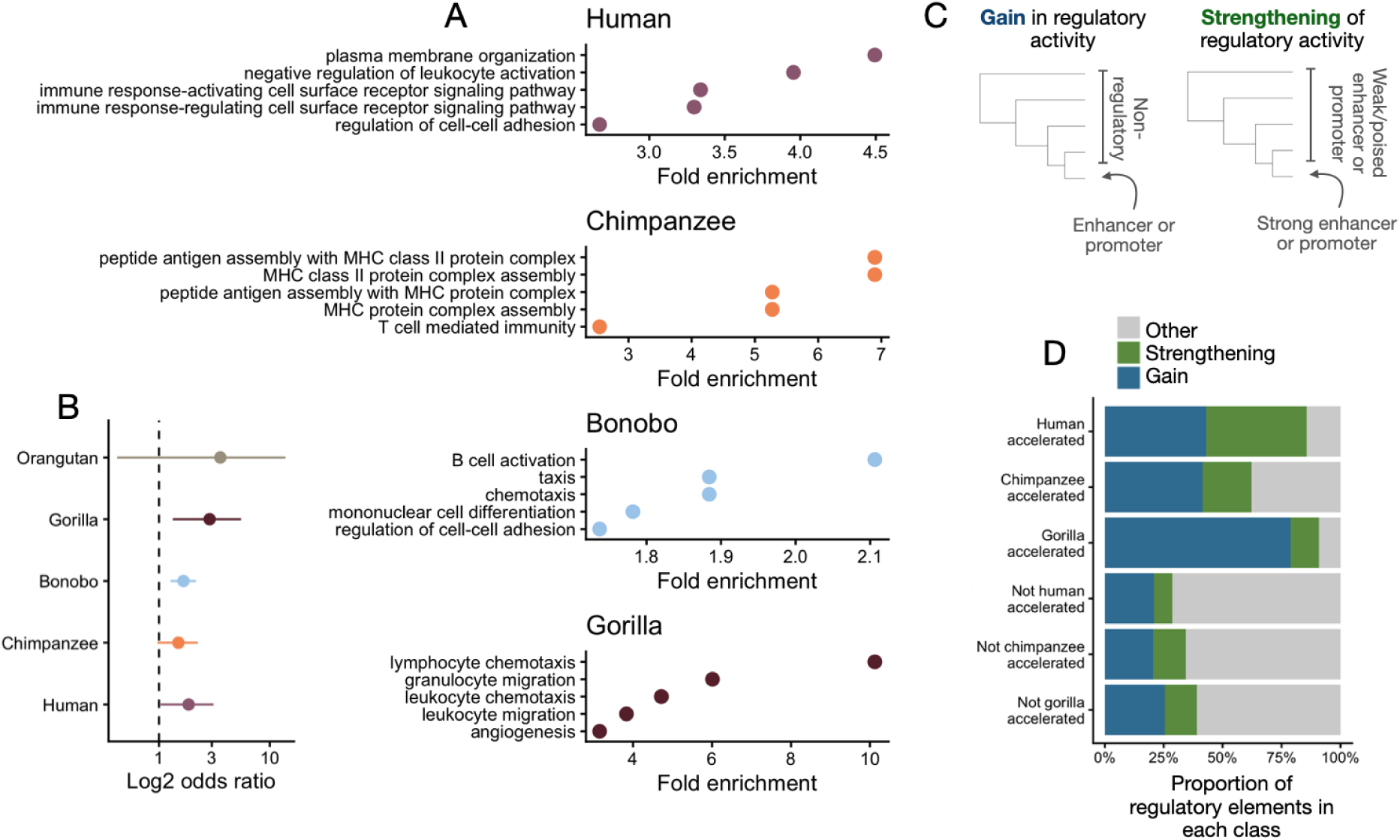
Biological properties of genes with accelerated evolution in terminal branches. A) Top 5 biological pathways enriched for genes accelerated in the focal lineage (gene ontology enrichment analysis, all adjusted p-values<0.05). B) Overlap between accelerated, lineage-specific sequence evolution [27] and accelerated, lineage-specific gene expression evolution. The x-axis point estimate represents the log_2_ odds ratio from a Fisher’s exact test, and the bars represent 95% confidence intervals. C) Definitions of regulatory elements that gained activity or strengthened their activity in the focal species relative to all other species on the tree (note that these annotations were performed separately for human, chimpanzee, and gorilla, but only one example focal species is shown in the heuristic). D) Proportion of regulatory elements with a species-specific state that gained activity, strengthened their activity, or adhered to some other pattern; barplots are stratified to compare regulatory elements associated with genes that did versus did not exhibit credible increases in gene expression in the focal species. For all plots, we define taxonomic groupings using common names: human (*Homo sapiens*), chimpanzee (*Pan troglodytes*), bonobo (*Pan paniscus*), gorilla (*Gorilla gorilla*), and orangutan (*Pongo abelii*).

Given that parasites and pathogens are also thought to drive selection in immune genes at the sequence level [59], we also tested whether accelerated gene expression evolution was accompanied by accelerated sequence evolution. To do so, we drew on a recently developed catalog of lineage-specific accelerated regions (LinARs) [27]. Specifically, Bi and colleagues used whole-genome sequences from 49 primate species to identify regions that have accumulated mutations at a faster rate on a particular branch relative to the rest of the tree. Using these LinAR annotations, we found significant overlap between sequence- and transcriptome-accelerated genes for humans (Fisher’s exact test: odds=1.85, p=0.025), bonobos (odds=1.66, p=2.43×10^−4^), and gorillas (odds=2.86, p=4.26×10^−3^), with non-significant effects in the expected direction for chimpanzees (odds=1.50, p=0.061) and orangutans (odds=3.58, p=0.114) (**Figure 3B**). These results suggest that gene expression evolution is generally accelerated in immune-related processes across apes, potentially consistent with previously observed Red Queen dynamics in primate immune systems [63]. Further, in many cases, lineage-specific acceleration at the transcriptomic level is likely driven by lineage-specific evolution at the sequence level.

### Accelerated gene expression evolution intersects with lineage-specific epigenomic variation

Mechanistically, lineage-specific gene expression evolution could be driven, at least in part, by changes in the epigenomic regulatory state associated with accelerated genes. For example, if a region of the genome that was previously non-regulatory gains enhancer or promoter function, this change could generate lineage-specific increases in gene expression levels (**Figure 3C**). To test this hypothesis, we drew on regulatory state annotations from a previous study of human, chimpanzee, gorilla, orangutan, and rhesus macaque LCLs [61]. We extracted previously assigned regulatory states for each species, which were derived from chromatin accessibility (ATAC-seq) and histone modification data (ChIP-seq of H3K4me1, H3K4me3, H3K36me3, H3K27ac, and H3K27me3) with the following set of states: ambiguous enhancer, ambiguous promoter, poised enhancer, poised promoter, strong enhancer, strong promoter, weak enhancer, weak promoter, or non-regulatory. For all analyses, we collapsed weak and poised categories into “weak/poised promoter” and “weak/poised enhancer” and excluded ambiguous annotations (see Methods). Focusing on humans, chimpanzees, and gorillas—the species included in [61] for which we also identified appreciable numbers of genes with credible increases in gene expression levels—we tested for overlap between lineage-specific regulatory states and patterns of gene expression evolution.

We found that human-accelerated, chimpanzee-accelerated, and gorilla-accelerated genes were all enriched near regulatory elements with distinct states in the focal species relative to all other species (e.g., for humans, elements where there was a different state in this species relative to chimpanzee, gorilla, orangutan, and macaque, who all shared the same state) (Fisher’s exact test, human: odds=2.15, p-value=0.010; chimpanzee: odds=1.50, p=0.012; gorilla: odds=2.87, p-value=3.97×10^−12^). For regions that fit this pattern, 43% (human), 42% (chimpanzee), and 79% (gorilla) involved novel gains in regulatory activity (transitions from a non-regulatory element to an enhancer or promotor in the focal species). Additionally, 43% (human), 21% (chimpanzee), and 9% (gorilla) of species-unique regions involved the strengthening of regulatory activity (transitions from a weak/poised enhancer or promoter to a strong enhancer or promoter in the focal species) (**Figure 3D**). By comparison, lineage-specific states near genes that do not exhibit signatures of accelerated gene expression evolution were less likely to indicate gains in regulatory activity, though results varied across species. Specifically, only 20-25% of these states reflected novel gains (Fisher’s exact test: human: odds=2.83, p-value=0.089; chimpanzee: odds=2.76, p=2.35×10^−3^; gorilla: odds=10.81, p-value=2.28×10^−9^), and only 8-14% reflected strengthening of regulatory activity (Fisher’s exact test: human: odds=8.76, p-value=6.86×10^−4^; chimpanzee: odds=1.61, p=0.283; gorilla: odds=0.87, p-value=1; **Figure 3D**, **Table S9**).

### Differentially expressed genes have accelerated rates of evolution

Genes that respond to specific environmental conditions may evolve under less pleiotropic constraint and be more disease-relevant than constitutively expressed genes [47,66]. If so, their gene expression levels may also evolve at different rates than genes that are less plastic. To test this prediction, we first identified environmentally responsive genes by testing for differential expression between treatment and control conditions across the five *in vitro* treatments included in our study (aldosterone, caffeine, glucose, tunicamycin, and acetaminophen) using a joint model across all species. The treatments varied in the breadth and strength of their effect on LCL expression (as observed previously in studies focused on human LCLs alone [48,51,52]). Specifically, we identified no response to glucose, but 1260 DE genes in response to caffeine, with the other three treatments falling somewhere in between (11 aldosterone-responsive genes, 58 tunicamycin-responsive genes, 61 acetaminophen-responsive genes, adjusted p-value < 0.05; **Figure 4A**, **Figure S6-7**, **Table S10**). However, even for treatments where we identified no or few DE genes, tests for functional enrichment among genes ranked by effect size [67] recovered biologically plausible pathways, consistent with a primary limitation of sample size. For example, genes upregulated by glucose were most clearly enriched for oxidative phosphorylation (adjusted p-value=5.10×10^−7^) and ATP synthesis coupled electron transport (adjusted p-value=1.94×10^−6^), while genes upregulated by tunicamycin were most clearly enriched for response to endoplasmic reticulum stress (adjusted p-value=4.51×10^−6^, **Figure 4B**, **Figure S8**, **Table S11**). Further, *IL1RN* and *DDIT3* were among the top 5 genes associated with glucose and tunicamycin response, respectively. *IL1RN* has documented associations with diabetes and glycemic traits, inflammatory regulation affecting insulin signaling, and metabolic pathways involved in glucose homeostasis [68,69]. Similarly, *DDIT3* is a classic stress-induced transcription factor that is strongly upregulated during prolonged or severe endoplasmic reticulum stress [70,71] (**Figure 4C**).

**Figure 4.**
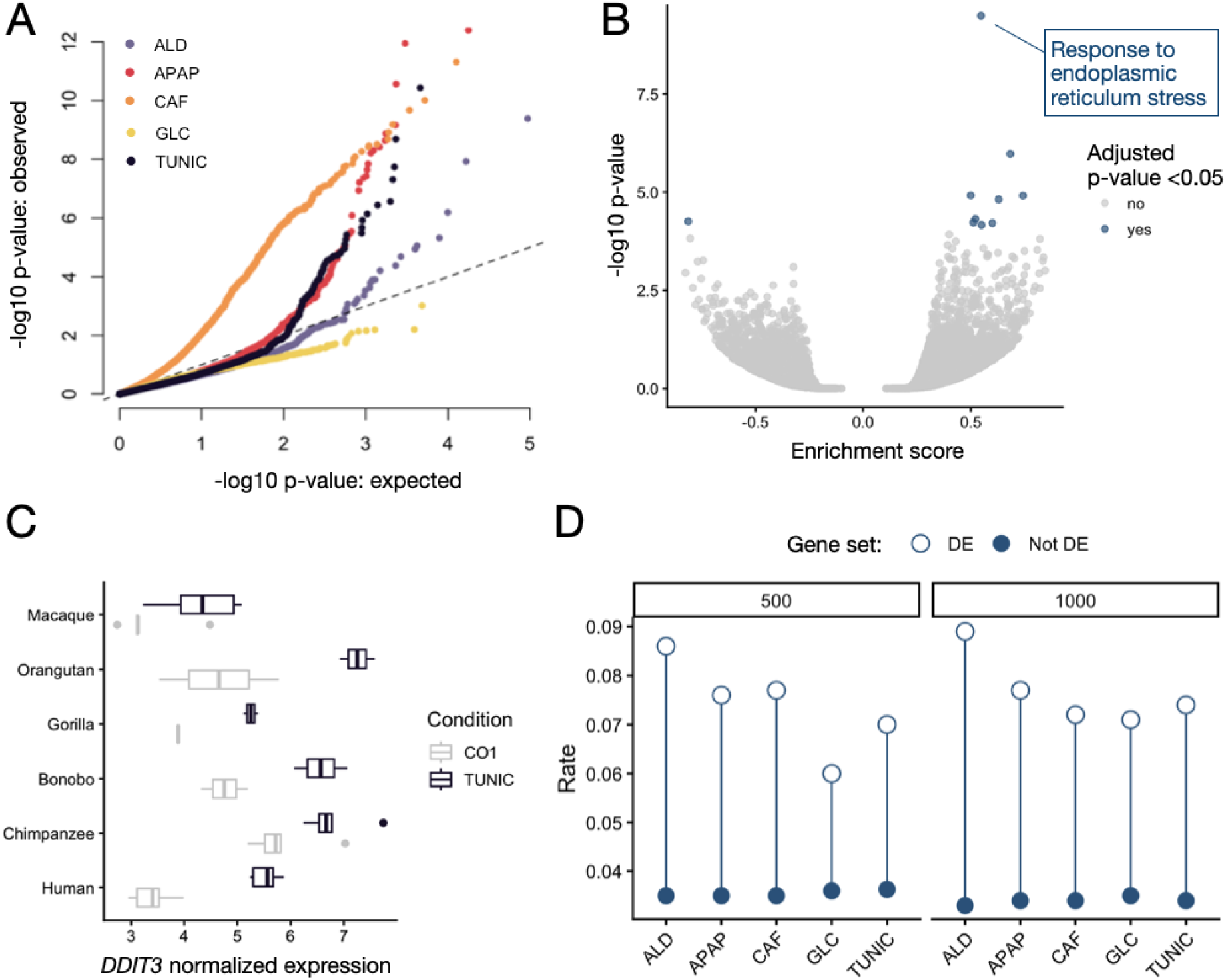
Accelerated gene expression evolution among differentially expressed genes. A) QQ-plot comparing the p-value distributions between an expected, uniform distribution and the observed values from differential expression analyses: aldosterone (ALD), acetaminophen (APAP), caffeine (CAF), glucose (GLC), and tunicamycin (TUNIC). The dotted line represents x=y. B) Volcano plot of biological pathways that exhibit enrichment for tunicamycin-responsive genes (gene set enrichment analysis, all adjusted p-values<0.05). Enrichment score was calculated by the clusterProfiler package in R and represents the degree to which a given biological pathway is overrepresented at the top or bottom of the ranked gene list. C) Boxplots of normalized *DDIT3* gene expression, as an example of a differentially expressed (DE) gene between the tunicamycin (TUNIC) and water condition (CO1). D) Evolutionary rates (y-axis) estimated for DE and not DE genes using the baseline water control expression data. Analyses were run comparing the top 500 or top 1000 DE genes, rank ordered by absolute effect size, versus all remaining genes. For all plots, we define taxonomic groupings using common names: human (*Homo sapiens*), chimpanzee (*Pan troglodytes*), bonobo (*Pan paniscus*), gorilla (*Gorilla gorilla*), orangutan (*Pongo abelii*), and macaque (*Macaca mulatta*).

Based on these results, and because we were interested in having similarly sized comparison sets across treatments, we rank-ordered each set of DE genes by their effect sizes and selected the top 500 or 1000 DE genes for each perturbation. We then used CAGEE to test whether the evolutionary rates of the top 500 or 1000 DE genes were accelerated compared to all other expressed genes, using expression values from the baseline water condition. Because our goal was to compare between environmentally-dependent and insensitive genes across the phylogeny as a whole, here we used a single rate model but allowed the rate to vary between DE genes and non-DE genes. We found that DE genes had, on average, a 2.1-fold higher rate of evolution (when using 500 genes) or a 2.3-fold higher rate (when using 1000 genes) than non-DE genes (**Figure 4D**). This result was consistent across all comparisons (**Table S12**).

We performed two secondary analyses to test the robustness of our result. First, we compared the top 500 (or 1000) to the bottom 500 (or 1000) DE genes (rank-ordered by their effect sizes) and found a 3.5-fold higher rate of evolution for the top DE genes compared to the bottom DE genes, suggesting our results are robust to the choice of background comparison (**Table S12**). Second, we performed a sliding window analysis where we estimated and compared the rates of evolution for the top 1-200, 201-400, 401-600, 601-800, 801-1000 DE genes (rank-ordered by their effect sizes) to the estimated rate for the bottom 1000 DE genes and observed similar evolutionary rate differences in all of these windows (**Figure S9**). Taken together, our results suggest that the faster rate of evolution we observe for immune-related gene expression—which has previously been attributed to the need for dynamism in response to pathogen threats [72]—likely generalizes to other classes of environmentally-dependent genes.

### Replication of qualitative patterns in primary immune cells

While LCLs have long served as a model for studies of genome, epigenome, and transcriptome evolution [60–62,73], they are immortalized through Epstein-Barr transformation and thus have biological features that are distinct from primary immune cells. Given this limitation, we used publicly available data from Hawash and colleagues [19], who collected gene expression data from primary immune cells exposed to bacterial lipopolysaccharide (LPS) and gardiquimod (GARD, a TLR7 agonist that mimics aspects of viral infection) challenges from humans, chimpanzees, olive baboons, and rhesus macaques (**Figure 5A**). Similar to our design, whole blood from each species was incubated with a given perturbation or a baseline control for 4 or 24 hours, after which white blood cells were purified and mRNA-seq was performed (**Figure 5B**). As expected, a PCA of the 151 samples from this study clustered by species rather than treatment, with principal component 1 separating the cercopithecoid monkeys (baboons and macaques) from the great apes (**Figure 5C**).

**Figure 5.**
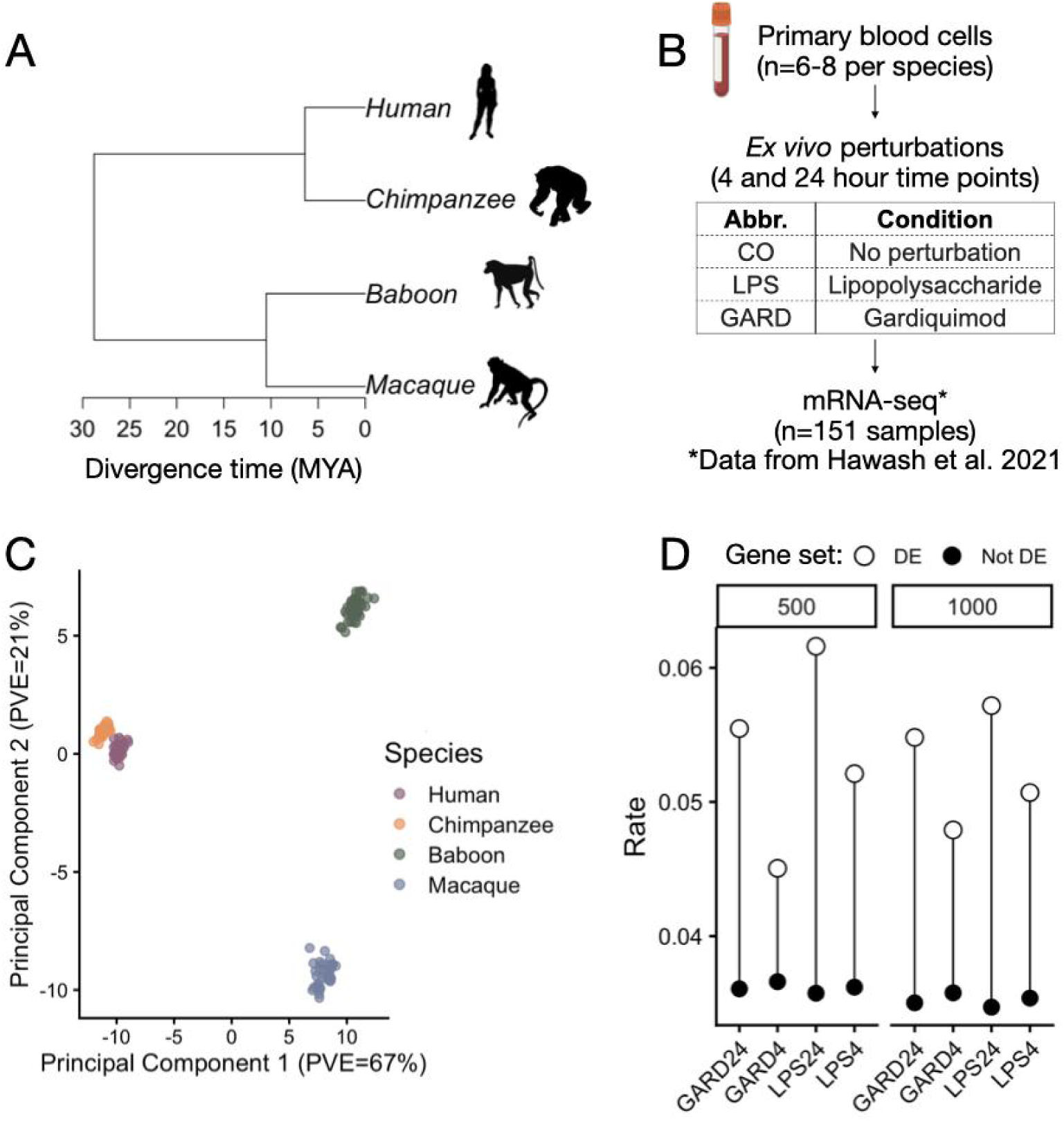
Replication of main results in primary immune cells. A) Phylogeny of species included in the Hawash et al. study, with divergence times from [74] and B) overview of experimental design. Whole blood from multiple species was exposed to baseline and treatment conditions—LPS and GARD—for 4 and 24 hours. After incubation, white blood cells were isolated for gene expression profiling. Evolutionary divergence times (million years ago, MYA) were obtained from [74]. Silhouette images were adapted from http://phylopic.org/ and courtesy of T. Michael Keesey, Tony Hisgett, and Gareth Monger (http://creativecommons.org/licenses/by/3.0/). Additional images were obtained from BioRender. C) PCA of white blood cell gene expression data across species and treatments. PVE = percent variance explained. D) Evolutionary rates (y-axis) estimated for differentially expressed (DE) and not DE genes using the control expression data. Analyses were run using the top 500 and top 1000 DE genes, rank ordered by absolute effect size. For all plots, we define taxonomic groupings using common names: human (*Homo sapiens*), chimpanzee (*Pan troglodytes*), baboon (*Papio anubis*), and macaque (*Macaca mulatta*).

When running CAGEE on the baseline condition for this dataset, we again found that a model with a higher rate of evolution on the chimpanzee branch fit the data best (Δ log-likelihood compared to single rate model: 233; **Table S6**). As in our original analyses, this result is unsurprising given the amplified power on short branches. To understand concordance between the datasets at the gene level, we asked whether similar genes were identified with credible increases or decreases in expression levels. As expected, we found substantial overlap between the set of genes with credible increases (Fisher’s exact test: odds=2.07, p-value<10^−16^) or decreases (odds=2.27, p-value<10^−16^) in gene expression identified in the two datasets (**Table S13**). Unlike for the LCL data set, gene ontology enrichment analysis of genes accelerated in specific lineages did not yield any significant pathways (at a 5% FDR threshold). However, among the larger set of genes accelerated in any lineage (n=1301), we again identified enrichment for immune-related processes including defense response to bacterium (adjusted p-value=2.49×10^−3^) and detection of stimulus (adjusted p-value=3.80×10^−3^) (**Figure S10**).

Finally, we asked whether genes differentially expressed in response to LPS or GARD (after 4 hours or 24 hours) also exhibited higher rates of evolution relative to the rest of the genome. Of note, all treatments produced strong responses with many DE genes detected, even at a more stringent 1% FDR threshold than we applied in the LCL data set (LPS 4 hours: 3630, LPS 24 hours: 3006, GARD 4 hours: 2186, GARD 24 hours: 3303). As expected, these DE genes were enriched for involvement in the innate immune response (LPS 4 hours: adjusted p-value=1.12×10^−13^), response to cytokine (LPS: adjusted p-value=2.01×10^−13^; GARD 4 hours: adjusted p-value=4.76×10^−11^), and defense response to virus (GARD: adjusted p-value=3.17×10^−13^) (**Figure S11**, **Table S14**). Using the same approach as our initial analysis, in which we rank-ordered genes by their effect sizes and selected the top 500 or 1000 DE genes for comparison, we again found an accelerated evolutionary rate for environmentally-dependent genes (**Figure 5D**, **Table S12**). On average across treatments, we observed a ∼1.5-fold increase in the rate of evolution for the top 500 DE genes, with the LPS 24 hour condition exhibiting the strongest acceleration. This result was consistent across treatments, time points, and the number of context-dependent genes included. Taken together, we therefore qualitatively replicate the evolutionary signal derived from our LCL dataset.

## Discussion

Understanding the evolution of gene expression in primates is important for addressing long-standing questions about the role of regulatory variation in adaptation [6], for identifying the drivers of between-species trait differences, and for gaining mechanistic insight into variation in disease susceptibility in humans [75–78]. In this study, we characterized the evolution of gene expression using Brownian motion models and data from the largest set of ape species to date [28,31,42,61]. Using immune cell gene expression data from LCLs and primary white blood cells, we found that genes with credible increases in expression levels are often concentrated in immune processes, overlap with lineage-specific accelerated regions inferred from DNA sequence data [27], and are enriched for species-specific shifts in epigenomic state. Because our study design included gene expression data collected at baseline and after challenge with diverse stimuli, we were also able to ask whether environmentally-dependent genes have accelerated evolutionary rates relative to environmentally-insensitive genes. We found that genes involved in environmental responses evolve at a consistently higher rate, consistent with theoretical expectations that genes with more narrow and context-dependent functions evolve under reduced constraint, potentially due to their reduced pleiotropy [30]. Given the increasingly appreciated relevance of context-dependent loci for disease-related traits in humans [47,50,66], our results suggest that context-dependent gene regulation may also be broadly important across primates—with accelerated evolution due to relaxed constraint and/or lineage-specific adaptation.

### Accelerated transcriptome evolution of environmentally-dependent genes

Our finding of accelerated evolution among environmentally-dependent genes represents a key advance over previous studies. Unlike prior comparative work, which has focused on either clarifying the evolutionary processes that shape baseline transcription [38] or identifying genes with differential responses to immune challenges between lineages without explicitly modeling gene expression evolution [19], we systematically tested for differences in evolutionary rates between gene sets using formal evolutionary models [36]. Our primary results qualitatively replicated in an independent dataset, suggesting generalizability across immune cell types, stimuli, and sample sizes. Moreover, they echo a larger body of cross-tissue comparative work [17,28,29,31,40], which has emphasized that genes expressed broadly across tissues tend to evolve more slowly than genes with tissue-specific expression patterns [30].

These results dovetail with ongoing discussions in human genomics about the importance of context-dependent regulatory variation for complex traits and disease. While two decades of GWAS have shown that non-coding variation plays a significant role in human disease [45,79,80], recent work suggests that genetic effects on gene expression—identified in the form of expression quantitative trait loci (eQTL)—exhibit genomic and evolutionary properties that differ from those of GWAS hits. Further, even in highly powered studies, eQTL show incomplete overlap with GWAS hits [46,47]. These observations have sparked interest in whether context-dependent eQTL—which are unmasked in certain tissues, environmental conditions, or life stages [47–51,81–84]—can help bridge the gap. Context-dependent eQTL often involve genes with critical developmental-, tissue-, or environment-specific functions [3,47,56] and may exhibit more “GWAS-like” properties, such as being further away from transcription start sites, more enriched in distal enhancers, and regulating selectively constrained, phenotypically-important genes [47,53,82,85]. For example, compared to eQTL consistently identified across conditions, diet-responsive eQTL identified in baboons [82] and tissue-specific eQTL identified in GTEx [45] were both more enriched for GWAS signals and occurred in more evolutionarily constrained, putatively disease-relevant genes. Theoretical arguments [86–88] suggest that this phenomenon may be driven by different evolutionary pressures on context-dependent versus non-context-dependent variants. Mutations that generate eQTL that are routinely expressed are consistently “visible” to selection and therefore can be rapidly purged if they are costly. In contrast, context-dependent eQTL may be “invisible” to selection in certain environments, tissues, or life stages, leading to weaker purifying selection, longer-term persistence, and greater contributions to disease. Thus, our results—that global gene expression levels recapitulate the known primate phylogeny, but that genes with context-dependent effects have accelerated evolutionary rates as compared to other genes—support a model in which gene expression evolution is dominated by negative and stabilizing selection [30,89] but suggest that genes with context-specific functions may accrue changes at faster rates than the genome as a whole.

### Accelerated gene expression evolution in immune pathways

In addition to providing a test of rate differences between environmentally-dependent and environmentally-insensitive genes, we also characterized general patterns of gene expression evolution in apes. While our power to identify genes with credible increases or decreases in expression was uneven across the tree, several results suggest that the genes we did identify within each lineage nonetheless reflect biological, rather than technical or statistical, patterns. For example, in the chimpanzee lineage, accelerated genes were most strongly enriched in MHC genes that bind antigens and display them for T-cell recognition, pointing to host-pathogen evolution and Red Queen dynamics as putative drivers of acceleration. The most strongly accelerated gene in chimpanzees was *B2M*, while the top accelerated *and* chimpanzee-differentially expressed gene was *HLA-C* (**Figure S12**). Both genes are part of the MHC class I system and, in humans, are tied to autoimmune disease risk [90–92] and infection outcomes, including HIV control [93,94]. For example, higher expression of HLA-C is associated with a lower viral load and slower HIV progression, presumably because it enables more effective cytotoxic T cell responses [93]. This is notable given the strong selective pressure that SIV is known to have exerted on the chimpanzee immune system [95], and the known differences in human-chimpanzee susceptibility to SIV/HIV and other diseases. Humans infected with HIV and hepatitis C develop stronger reactions and more severe complications than chimpanzees, for instance, and humans also appear to suffer more from autoimmune diseases such as asthma, psoriasis, and rheumatoid arthritis [78].

More broadly, genes with credible increases in expression levels in chimpanzees, bonobos, gorillas, and humans were all enriched for immune processes, though the individual pathways we detected varied between species. The concentration of expression increases in immune genes may arise from their fundamental role in responding to the environment (making them common in context-dependent gene sets [19]), and their importance in adaptive evolution in humans and other primates [59,79]. In agreement, we also observed concordance between sequence acceleration and transcriptome acceleration across species (similar to [96]), suggesting that lineage-specific regulatory changes often underlie transcriptional divergence.

### Limitations and future directions

Our study has several limitations. First, while some of the LCL perturbations are relevant to naturally occurring physiological or cellular processes (aldosterone, glucose, tunicamycin), others (acetaminophen, caffeine) are not molecules that non-human species naturally encounter, raising questions about how evolution could act on the transcriptional dynamics of these genes. We instead suspect that these genes are also sensitive to broader classes of molecules, such as hormonal, nutrient, or stress signaling molecules. Future work testing a larger panel of environmental perturbations, including additional ecologically and evolutionarily relevant perturbations, would be fruitful for confirming the patterns observed here. Second, our study focused on LCLs, which continue to be a useful model for gene expression evolution (Supplementary Text) but represent only one cell type. Expanding future work to additional cell types will be critical for confirming the patterns we observed here. Third, we did not explicitly test for selection with a stochastic process, such as Ornstein-Uhlenbeck (OU). Prior work has shown that OU models can have problems controlling the false positive rate [97]; thus, we focused on estimating evolutionary rates in a Brownian motion model. Future work could apply OU models with simulation-based null calibration to differentiate between fast evolution due to loss of constraint, versus fast evolution due to lineage-specific selection. Fourth, while we replicate the qualitative signal of faster gene expression evolution in context-dependent genes, the effect sizes are lower in our replication dataset. This effect size attenuation could reflect a winner’s curse effect [98,99] or result from the sparser phylogenetic tree in the replication data set. Finally, due to our modest sample sizes per species, we are underpowered to identify many differentially expressed genes, leading us to focus our rate difference tests on the top 500 or 1000 DE genes in response to each treatment. We note that imperfect classification of DE and non-DE genes should lead to an underestimation of evolutionary differences, rather than generating false positive results. Future comparative studies with larger sample sizes, including those with sufficient power to map context-dependent eQTL across species, will be important for clarifying the evolutionary and mechanistic basis of context-dependent gene expression [22].

## Methods

### Cell Lines

We worked with 26 previously established lymphoblastoid cell lines (LCLs) derived from humans (n=9), chimpanzees (n=5), bonobos (n=2), gorillas (n=2), orangutans (n=2), and rhesus macaques (n=5) (**Table S1**). Human LCLs derived from the 1000 Genomes Project were purchased from Coriell [100], nonhuman ape LCLs were established at the Max Planck Institute for Evolutionary Anthropology (MPI-EVA) from captive apes in zoo and sanctuary settings (see Supplementary Text), and macaque LCLs described in previous work [101] were shared with us by Yoav Gilad. Sex was balanced across each species.

### Cell Culture Experiments

LCLs were maintained at concentrations of 0.5-1.0 × 10^6^ cells/mL in RPMI (72400054, Gibco) supplemented with 10-20% fetal bovine serum (FB5001, Thomas Scientific) and 1% Penicillin/Streptomycin (30-002-CI, Corning), and cultured at 37°C in 5% CO_2_ and atmospheric O_2_. LCLs were cultured in parallel until they reached target densities and viabilities, at which time they were seeded in 24-well culture dishes with 0.5 × 10^6^ cells and 750 uL of media per well. After an overnight incubation period, control and perturbation treatments (**Table S2**) were added to the medium, with each condition isolated to a separate cell culture well. After a 4-hour exposure period, genome-wide gene expression patterns were measured at bulk resolution.

Given the difficulties of obtaining permits to share nonhuman primate cell lines between countries, human and rhesus macaque LCL experiments were performed at Vanderbilt University, and nonhuman ape LCL experiments were performed at MPI-EVA. All experiments used identical protocols. While performing experiments across two institutes likely generated some technical variation, our combined dataset shows expected evolutionary patterns (e.g., correlations between genomic and transcriptomic distance) and our main analyses focus on genes that are differentially expressed across all species.

### RNA Extraction and Sequencing

Total RNA was extracted from each sample using the Zymo Quick-RNA 96 kit (R1053, Zymo), following the manufacturer’s protocol. Strand-specific RNA-seq libraries were generated using the NEBNext Ultra II Directional RNA Library Prep Kit (E7760, NEB) with the NEBNext Poly(A) mRNA Magnetic Isolation Module (E7490, NEB). All libraries (n=169) were sequenced on the Illumina NovaSeq platform, with each sample sequenced to a minimum of 10 million uniquely mapped reads that were also assigned to a feature (i.e., annotated gene/transcript; **Table S3**). Library preparation and sequencing services were provided by GENEWIZ from Azenta Life Sciences (Leipzig, Germany) and the VANTAGE Sequencing Core at Vanderbilt University.

### Low level data processing

We obtained the genome builds hg38, panTro6, panPan3, gorGor6, ponAbe3, and rheMac10 and the chain files hg38ToPanTro6.over.chain.gz, hg38ToPanPan3.over.chain.gz, hg38ToGorGor6.over.chain.gz, hg38ToPonAbe3.over.chain.gz, and hg38ToRheMac10.over.chain.gz from the UCSC genome browser [102] and used custom scripts to construct a multi-species consensus genome across all 6 species (as described in [103] with scripts available at https://github.com/kennethabarr/ConsensusGenomeTools [104]). Gene annotations for this multi-species consensus reference were based on the hg38 RefSeq annotation downloaded from the UCSC genome browser [102]. Sequenced reads were mapped to the multi-species consensus reference using STAR (version 2.7.7a) [105], and gene expression levels were quantified using the featureCounts function in Subread (version 2.1.1) using standard parameters [106]. All downstream processing and analysis steps were performed in R (v4.4.0) unless otherwise stated.

Read counts were filtered for autosomal, protein coding genes using Ensembl annotations and normalized with the weighted trimmed mean of M-values (TMM) option in edgeR [107]. We next filtered for genes with median, normalized expression levels > 1 within at least one species, which resulted in 12,455 total genes. Focusing on these genes, we merged the original per-species counts matrices to generate a single multi-species matrix, which we renormalized again using the weighted TMM option in edgeR [107]. We applied the prcomp function for principal components analysis and the Mantel test function in the vegan R package to compare pairwise genetic distance to pairwise mean expression distance [108]. We used the ape R package to visualize species trees [109] and the hclust function to perform hierarchical clustering on the mean, per-species expression values. For the main analyses focused on the LCL dataset, we used divergence times from [64,65]; for the replication analyses focused on the white blood cell dataset, we used divergence times from [74] (**Table S4**).

### Differential expression analyses

We performed differential expression analyses using limma from the R package voom [110]. For each treatment—aldosterone, caffeine, glucose, tunicamycin, and acetaminophen—we ran a linear model comparing treated to control samples (using the water vehicle control for glucose and caffeine, and the ethanol vehicle control for aldosterone, tunicamycin, and acetaminophen). All models controlled for species and read depth (number of reads mapped to autosomal, protein coding genes). We performed multiple hypothesis testing correction using a Benjamini Hochberg false discovery rate approach implemented in p.adjust [111].

To understand the biological relevance of treatment-responsive genes, we used gene set enrichment analysis implemented in the R package clusterProfiler [67]. We rank ordered genes by their effect size, and tested for enrichment within known biological processes (5% FDR). We also repeated the above analyses (differential expression and gene set enrichment) comparing the water-treated to the ethanol-treated samples. These analyses did not reveal any significantly DE genes (exploring up to a 20% FDR) nor did they uncover any significantly enriched biological processes.

### Simulations to evaluate statistical power

We used CAGEE v1.2 [36,37] to simulate gene expression across the 6 taxa using the species tree in Figure 1A (see also **Table S4**). We simulated data for 12,468 genes using a baseline evolutionary rate of 0.0001. We also tested two-rate models where the rate was multiplied by a factor in a specific lineage. We varied the factor across 0.5, 0.75, 1 (the null model), 1.25, 1.5, 1.75, 2, 3, 4, and 5. We varied the lineage specific rate across H, C, B, G, O, M, CB, HCB, HCBG, HCBGO. We ran CAGEE on the simulated data and evaluated the power and false positive rates across 100 replicates for each parameter combination. In all cases, we applied a likelihood ratio test comparing the alternative model to the null model of a single rate across the entire tree.

Our null simulations revealed an inflated false positive rate across all alternative models when using the critical value for a chi-square test with one degree of freedom (**Figure S2**). Consequently, we set the likelihood ratio test critical value to the empirically determined value for each clade that resulted in a false positive rate of 5% for all downstream analyses.

### CAGEE analysis

We used CAGEE v1.2 [36,37] to study gene expression evolution. We computed the mean gene expression levels using counts per million normalized expression (DEseq2 [112]) in each baseline condition (water and ethanol) separately across all individuals within each species. We ran CAGEE using the bounded Brownian motion model using the species phylogeny in Figure 1A (see also **Table S4**).

We tested 12 nested models where the evolutionary rate was allowed to vary across different parts of the phylogeny. Specifically, we evaluated: 1) a base model of a single evolutionary rate across the entire tree, 2) six two-rate models where a single lineage was allowed its own rate and the rest of the tree was allowed a different rate, 3) four two-rate models where multiple species with shared ancestry were allowed to share a rate and the rest of the tree had a different rate, and 4) a model where all nodes of the tree, internal and external, were allowed their own rate. We used the negative log-likelihood values computed by CAGEE to compare models and considered significant differences as those with a log-likelihood difference greater than the 5% false positive rate cutoff from our null simulations.

To study the evolution of DE genes, we ranked genes by the absolute effect size in the differential gene expression analysis for each of the five treatments. Then, we annotated the top 500 DE genes in one group and the rest of the genes in a second group using CAGEE sample type annotations. We ran CAGEE on the baseline gene expression data and used the single rate tree to avoid over-parameterization. We allowed CAGEE to estimate one evolutionary rate for the DE genes and one for the rest of the genes. We carried out four additional similar analyses. First, we repeated the analysis using the top 1000 genes. Second, we repeated the analysis using the top 500 genes and the bottom 500 genes ranked by absolute value of DE effect size. Third, we repeated the analysis using the top 1000 genes and the bottom 1000 genes ranked by absolute value of DE effect size. Lastly, for the LCL data set, we performed a sliding window analysis in which we estimated the rates of evolution for the top 1-200, 201-400, 401-600, 601-800, 801-1000 genes ranked by absolute value of DE effect size and compared these to the estimated rate of evolution for the same set of bottom 1000 genes with the smallest absolute value of DE effect size.

### Biological annotation of accelerated genes

Along with evolutionary rates, CAGEE also outputs a list of genes with significant changes in evolutionary rate across the phylogeny. We extracted the genes identified as having significant acceleration on each terminal branch. For all species except macaque, for which no significantly accelerated genes were identified, we used the per-species gene sets as the input for gene ontology enrichment analyses implemented in the R package clusterProfiler (5% FDR) [67]. All analyses compared the focal set of significant genes to the background set of all expressed genes included in our analyses.

To compare the genes with significantly accelerated rates of gene expression evolution to genomic regions with significantly accelerated sequence evolution, we downloaded annotations for lineage-specific accelerated regions (LinARs) from [27]. For each species, we used Fisher’s Exact Tests to test whether genes that exhibited an accelerated rate of gene expression evolution in our analysis increased the odds of being located within or overlapping LinARs (i.e., regions identified as fast-evolving on the branch leading to the same species at the sequence level).

We used a similar approach to compare the genes that exhibited significantly accelerated rates of gene expression evolution in chimpanzees to properties captured in other datasets. First, we used Fisher’s exact tests to ask whether genes identified as differentially expressed between chimpanzees and macaques or chimpanzees and humans in a previous study of LCLs [42] were enriched among the chimpanzee-accelerated genes identified in this study. For this comparison, we used the definitions of differential expression from the original paper (1% FDR) and restricted our comparison to genes included in both our study and the comparison study.

Second, we downloaded regulatory annotations from a previous study of epigenetic marks in human, chimpanzee, gorilla, Sumatran orangutan, and rhesus macaque LCLs [61]. This study applied chromHMM to chromatin accessibility (ATAC-seq) and histone modification data (ChIP-seq of H3K4me1, H3K4me3, H3K36me3, H3K27ac, and H3K27me3) generated in parallel for each species to assign regulatory states to 200 bp genomic windows. We focused on orthologous regulatory elements provided by the authors in Supplementary Table 3. Each regulatory element was preannotated as associated with a given gene based on proximity and 3D chromatin maps in human LCLs (GM12878), which provide information about physical chromatin interactions. A total of 28703 orthologous regulatory elements were provided, 20339 of which were retained for downstream analyses because they could be assigned to a single (unique) gene. Each element was annotated with one of the following state possibilities for each species: ambiguous enhancer, ambiguous promoter, poised enhancer, poised promoter, strong enhancer, strong promoter, weak enhancer, weak promoter, or non-regulatory. We combined the poised and weak categories to make a “poised/weak promoter” and a “poised/weak enhancer” category. Using these definitions, 22% of elements had identical regulatory states across all species. In contrast, 7% of elements had a distinct state in humans relative to all other species (who shared the same state); the percentages for parallel analyses constructed with chimpanzees and gorillas as the focal species were 3% and 4%, respectively. After filtering for regulatory elements associated with genes expressed in our LCL dataset, we retained 13072 orthologous regions.

Following filtering, we retained 335, 254, and 340 elements where the human, chimpanzee, or gorilla state differed from all other species. When deriving these lineage-specific regulatory states and in subsequent analyses, we excluded regions where the regulatory state was labeled as ambiguous in the focal species. We then asked whether each set of lineage-specific elements was enriched near human-chimpanzee-, or gorilla-accelerated genes, respectively, using a Fisher’s exact test. To compare the patterns we observed near chimpanzee-accelerated genes to background expectations, we repeated the above analyses for the 645 elements with regulatory state differences between chimpanzees and the rest of the species that were not near chimpanzee-accelerated genes, using a Fisher’s exact test.

### Replication analyses in primary cells

We downloaded sample-specific processed counts files (output from Kallisto [113]) from the Sequencing Read Archive for accession number GSE155918 [19]. We combined the counts files across samples and species based on Ensembl gene identifiers. As in our main analyses, we again focused on autosomal, protein coding genes, normalized the counts within each species using the TMM option in edgeR [107], filtered for genes with normalized expression levels > 1 within at least one species, and then merged and renormalized the multi-species matrix. This approach left us with 8971 filtered genes for analysis. We repeated the pipelines described above for principal components analysis, differential expression testing, CAGEE, and gene ontology enrichment analysis of significantly accelerated genes. To compare the genes identified in both datasets as significantly accelerated or decelerated in any terminal branch, we used Fisher’s exact tests subsetting to the set of genes common to both the LCL and white blood cell datasets.

## Acknowledgments

We thank Kelly Williams, Komal Kadyan, Jenny Busch, and Michelle Roderer for their assistance with the cell culture experiments. We thank Anne Fischer, Michel Halbwax, and members of the Chimpanzee Sanctuary & Wildlife Conservation Trust, Tacugama Chimpanzee Sanctuary, and the Leipzig Zoo Wolfgang Köhler Primate Research Centre for sharing biological samples to generate LCLs. We thank Svante Pääbo for establishing and sharing these LCLs, as well as Chris Tyler-Smith for sharing a gorilla LCL. We thank members of the Lea and Housman labs for their feedback on this work.

## Code and data availability

All FASTQ files are available on the NCBI’s Gene Expression Omnibus under accession numbers GSE334802 and GSE334804. All code is available at https://github.com/durvasula-lab/evolution_response_primates

