## Supplementary Text and Figures for "Accelerated Evolution of Context-Dependent Gene Expression in Primate Immune Cells"

**This file includes:**


- Supplementary Text
- Figure S1. Variation in LCL gene expression data
- Figure S2. Distribution of likelihood ratio test statistics in null simulations
- Figure S3. Number of genes with credible expression increases in a given lineage
- Figure S4. Overlap between genes with credible increases or decreases in expression in a given lineage
- Figure S5. Gene ontology results for genes with credible increases or decreases in expression in a given lineage
- Figure S6. Differentially expressed genes in response to perturbations in LCLs
- Figure S7. Overlap between differentially expressed gene sets in LCLs
- Figure S8. Gene ontology analysis of differentially expressed genes in LCLs
- Figure S9. Results from sliding window analyses
- Figure S10. Gene ontology analysis of significantly accelerated genes in white blood cells
- Figure S11. Treatment responsive genes in white blood cells
- Figure S12. Acceleration and differential expression of MHC genes in chimpanzees
- Supplementary References


**The following tables are provided in separate file(s):**


- Table S1. Lymphoblastoid cell line information
- Table S2. Treatment information
- Table S3. Experiment, sample, and sequencing metadata
- Table S4. Newick formatted trees that were used in the CAGEE analyses
- Table S5. Power to detect evolutionary rate changes in simulated data
- Table S6. Log-likelihoods of all evolutionary models tested
- Table S7. Genes with credible increases in gene expression levels in LCLs
- Table S8. Gene ontology of genes with credible changes in gene expression levels in LCLs
- Table S9. Regulatory elements with unique states in chimpanzee, gorilla, or human
- Table S10. Genes significantly differentially expressed in response to each treatment
- Table S11. Gene set enrichment analysis of the response to each treatment
- Table S12. Evolutionary rate comparisons between differentially expressed (DE) and not DE genes (in baseline data)
- Table S13. Genes with significantly accelerated evolutionary rates in white blood cells
- Table S14. Gene ontology results from differentially expressed genes in white blood cells

### Supplementary Text

*Lymphoblastoid cell lines (LCLs) as a model for gene expression evolution*

Robust comparative studies of genome function, especially across diverse cell states, requires access to large numbers of live cells from many individuals and species, but ethical, legal, and practical limitations often restrict access to such biological materials. This limitation is especially true for non-human great ape species, which are the most relevant group for understanding the evolution of human-specific traits. To overcome this issue, many researchers have used lymphoblastoid cell lines (LCLs), which are immortalized from primary immune tissue (B cells) and can thus be kept long-term and grown at scale.

In humans, LCLs have been used extensively for functional genomic work, and previous studies have shown that 1) genomic results from LCLs replicate in primary tissues and 2) gene expression levels in newly established LCLs maintain a strong individual signature [1–5]. However, gene regulation in LCLs is not identical to their progenitor B cells, and the transformation process is known to induce certain artifacts [6,7]. Nonetheless, across species, LCLs have long-served as a workhorse for comparative functional genomic insight [8–10], with strong evidence that they are an appropriate model for studying generalizable evolutionary processes [9]. Additionally, the clonal nature of LCLs provides a cell type-specific system, minimizing confounding factors associated with cell type heterogeneity. Thus, while a handful of previous studies have combined comparative functional genomic approaches in primary tissues with *ex vivo* perturbations [11,12], work in this area has been limited in terms of both species diversity and perturbation diversity [13]) because gaining access to the necessary biological materials from non-human great ape species is practically very difficult.

*Cell Line Generation, Acquisition, and Maintenance*

The LCLs used in this study were previously generated from peripheral blood mononuclear cells (PBMCs) collected from human and nonhuman primates in compliance with applicable national and international regulations.

Human LCLs were previously generated from individuals included in the 1000 Genomes study [14] and ordered from the Coriell Institute. Macaque LCLs (described in [15]) were previously generated in Dr. Yoav Gilad's group and shared with the Lea lab. All chimpanzee LCLs were previously generated in Dr. Svante Pääbo's lab from animals housed at the Chimpanzee Sanctuary and Wildlife Conservation Trust and the Tacugama Chimpanzee Sanctuary. Blood samples were opportunistically collected by Michel Halbwax and Anne Fischer in 2007 and 2008 during routine health assessments [16]. One gorilla LCL was previously shared by Dr. Chris Tyler-Smith at the University of Oxford CRC Chromosome Molecular Biology Group with Dr. Pääbo's lab. All other nonhuman ape LCLs (2 bonobo, 1 gorilla, 2 orangutan) were previously generated in Dr. Pääbo's lab from captive animals housed at the Leipzig Zoo Wolfgang Köhler Primate Research Centre. Blood samples were drawn during routine medical exams conducted by veterinarians between 2001 and 2009, and only surplus samples were used.

For all LCLs, PBMCs were isolated from whole blood samples using a Ficoll gradient, and B-cells were transformed and immortalized through exposure to the Epstein Barr Virus derived from the B95-8 EBV-producing marmoset B-cell line. Established LCLs were maintained in RPMI (72400054, Gibco) supplemented with 10-20% fetal bovine serum (FB5001, Thomas Scientific) and 1% Penicillin/Streptomycin (30-002-CI, Corning). Cells were cultured at 37˚C in 5% $CO_2$ and atmospheric $O_2$ and maintained at concentrations of 0.5-1.0 x $10^6$ cells/mL.

#### Supplementary Figures

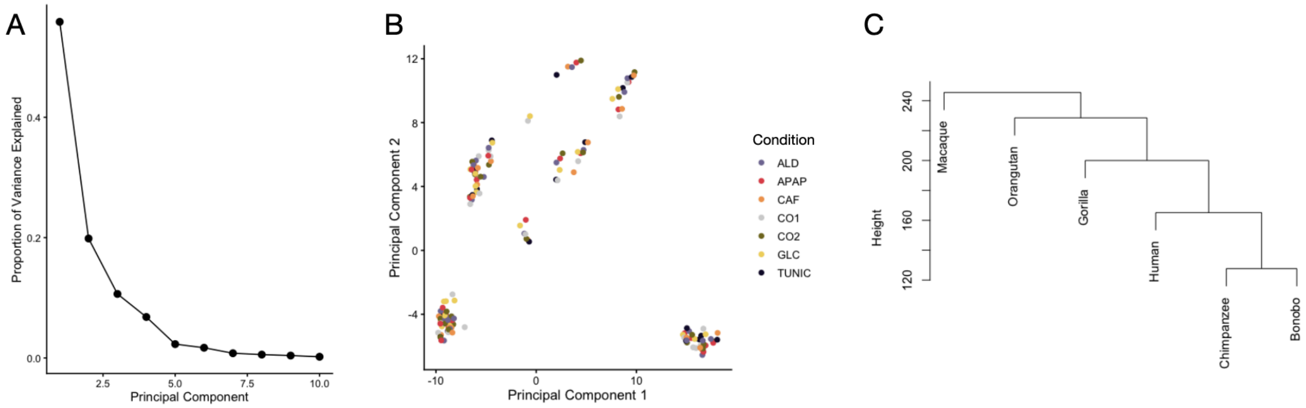


**Figure S1. Variation in LCL gene expression data.** A) Scree plot showing the proportion of variance explained by each PCA for the PCA of LCL gene expression data across species and conditions. B) Principal component 1 and 2 colored by condition rather than species (as in Figure 1). C) Hierarchical clustering of LCL gene expression data, focusing on a matrix of mean normalized expression levels by species. Treatment acronyms: ALD (aldosterone), APAP (acetaminophen), CAF (caffeine), CO1 (water), CO2 (ethanol), GLC (glucose), TUNIC (tunicamycin).

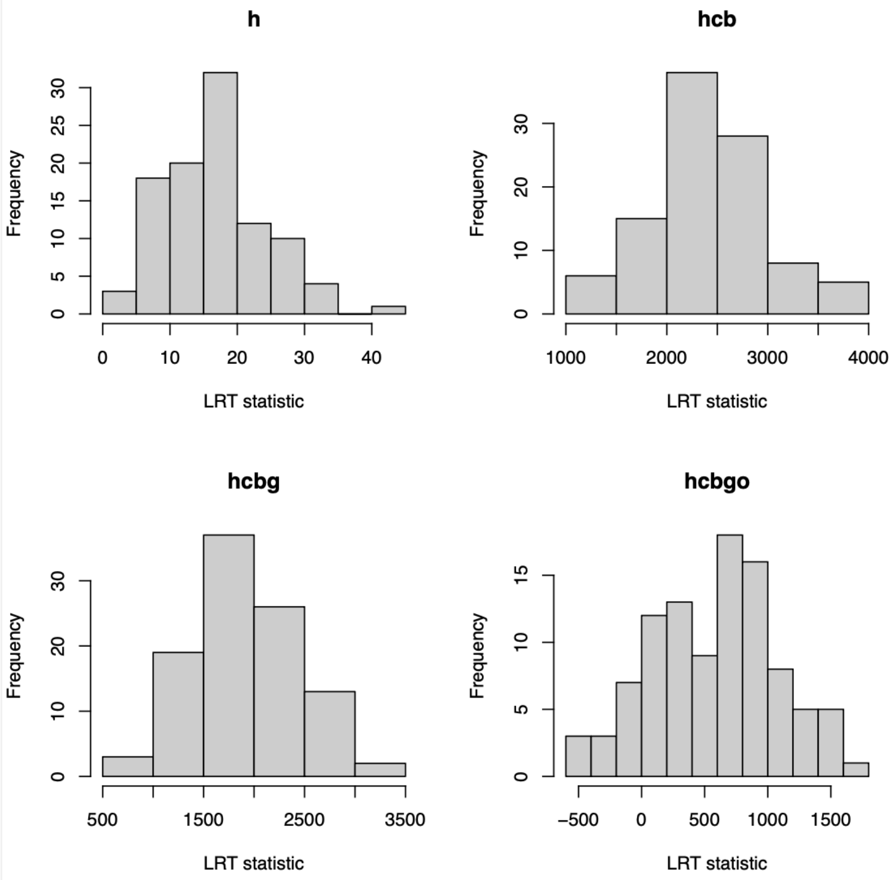


**Figure S2. Distribution of likelihood ratio test statistics in null simulations.** For each panel, we show the result of applying a likelihood ratio test to simulated data where there is no change in the rate of gene expression evolution. Then, we run CAGEE to infer the evolutionary rate with one rate for a focal branch (annotated in the plot title: h (human lineage), hcb (human-chimpanzee-bonobo lineage), hcbg (human-chimpanzee-bonobo-gorilla lineage), hcbgo (lineage of all great apes)) and another rate for the rest of the tree. The likelihood ratio test compares the two-rate model to the single rate model. In all cases, the distribution of test statistics is inflated above a chi-squared distribution with one degree of freedom. In the main text, we adjusted the likelihood ratio test significance threshold to control for a 5% false positive rate.

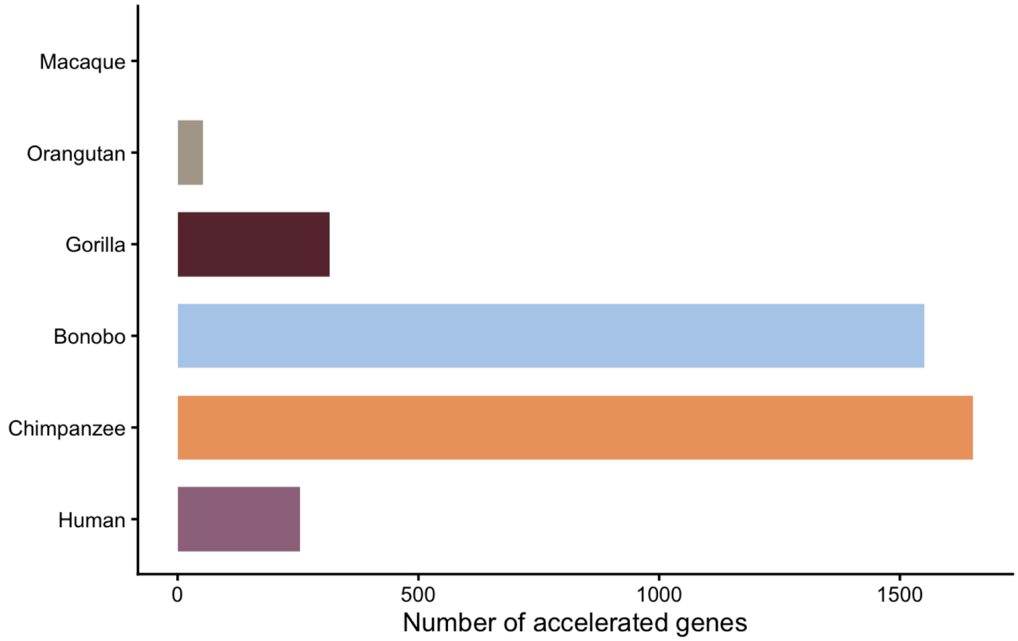


**Figure S3. Number of genes with credible expression increases in a given lineage.** Number of genes for which CAGEE identified a credible expression increase on that branch relative to the immediate parent node. We note that these numbers are strongly driven by power differences inherent in the tree structure, but we provide them as context for downstream biological analyses.

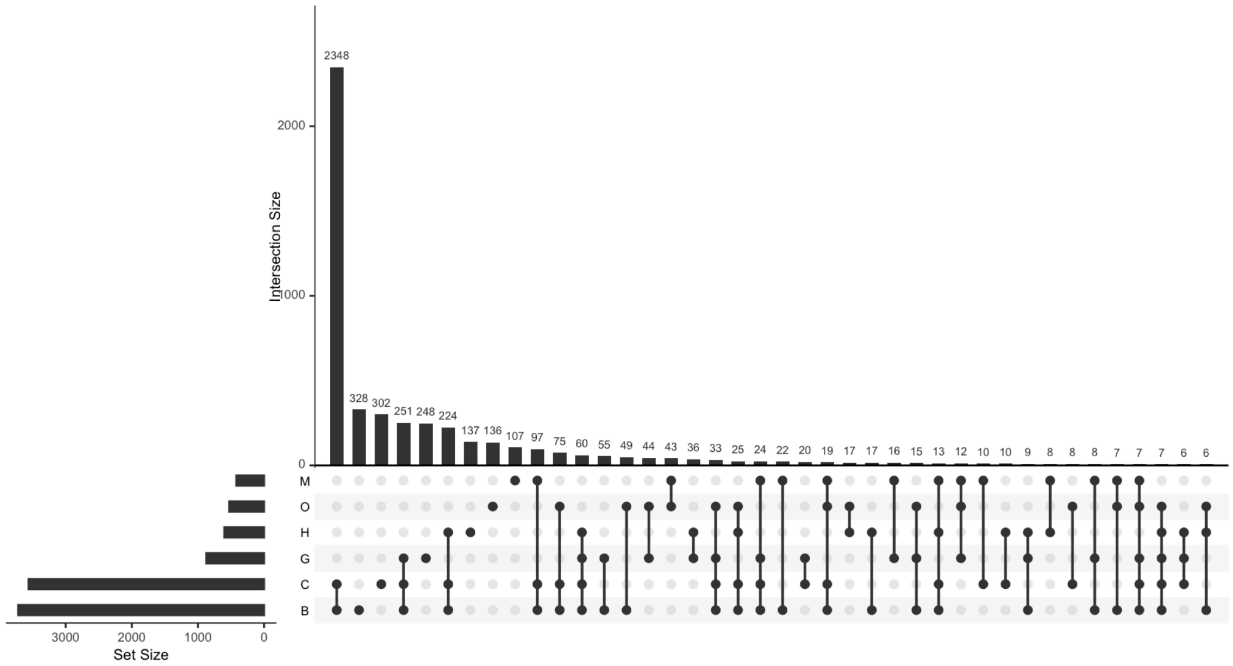

**Figure S4. Overlap between genes with credible increases or decreases in expression in a given lineage.** Upset plot shows the overlap in gene identities for all genes for which CAGEE identified a credible expression difference on that branch relative to the immediate parent node. H = human (*Homo sapiens*), C = chimpanzee (*Pan troglodytes*), B = bonobo (*Pan paniscus*), G = gorilla (*Gorilla gorilla*), O = orangutan (*Pongo abelii*), and M = macaque (*Macaca mulatta*).

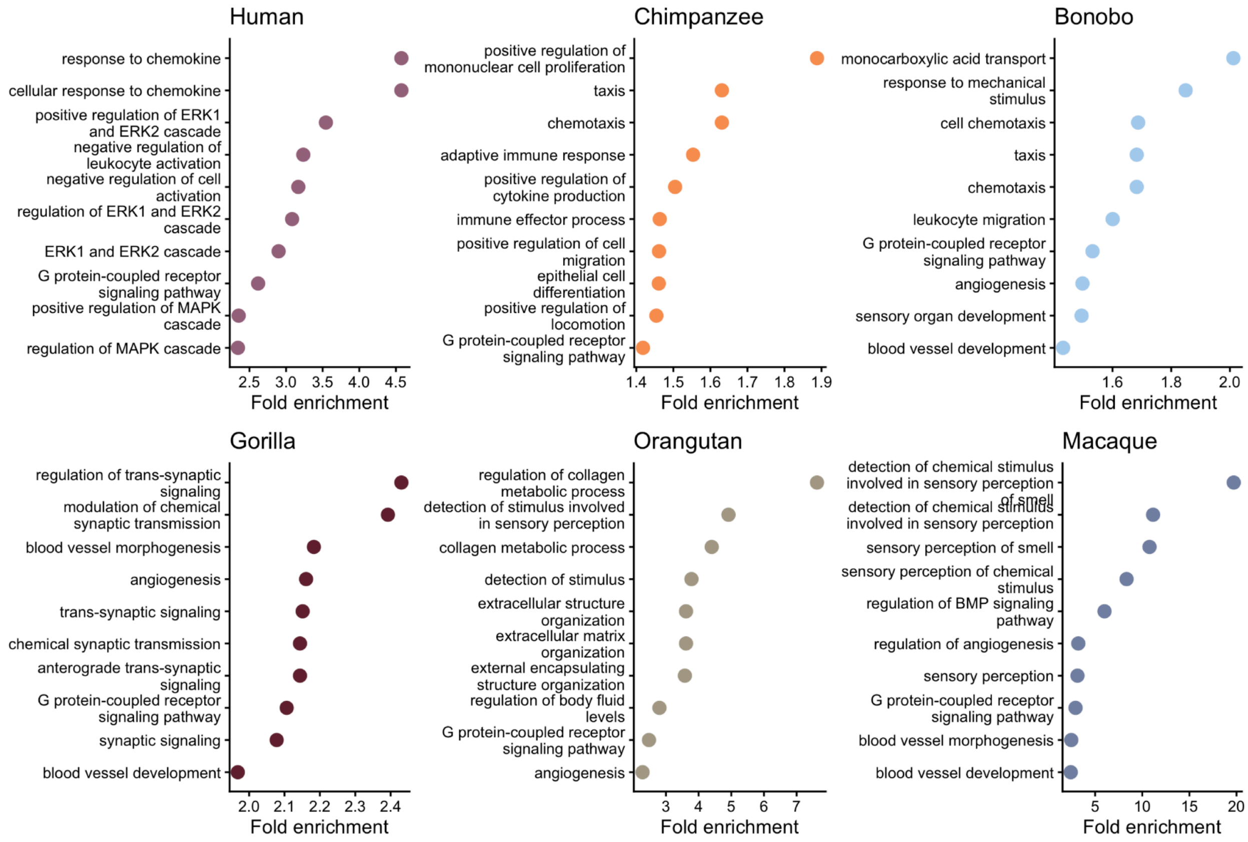


**Figure S5. Gene ontology results for genes with credible increases or decreases in expression in a given lineage.** The top 10 biological pathways identified via gene ontology enrichment analysis are shown for each focal lineage (all adjusted p-values<0.05).

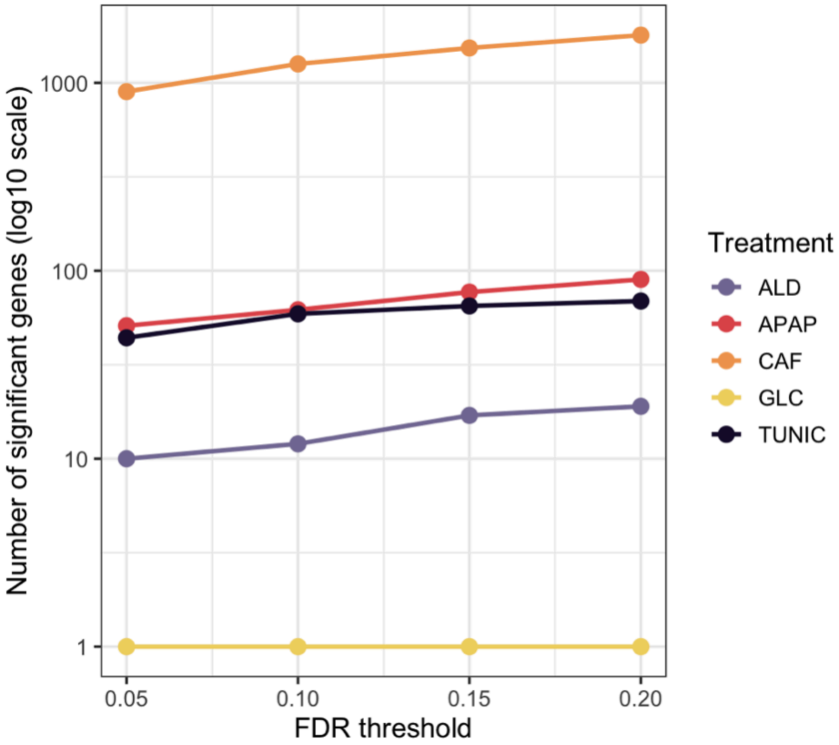


**Figure S6. Differentially expressed genes in response to perturbations in LCLs.** The number of significantly differentially expressed genes between treatment and control conditions (y-axis) across a range of FDR thresholds (x-axis). Treatment acronyms: ALD (aldosterone), APAP (acetaminophen), CAF (caffeine), GLC (glucose), TUNIC (tunicamycin).

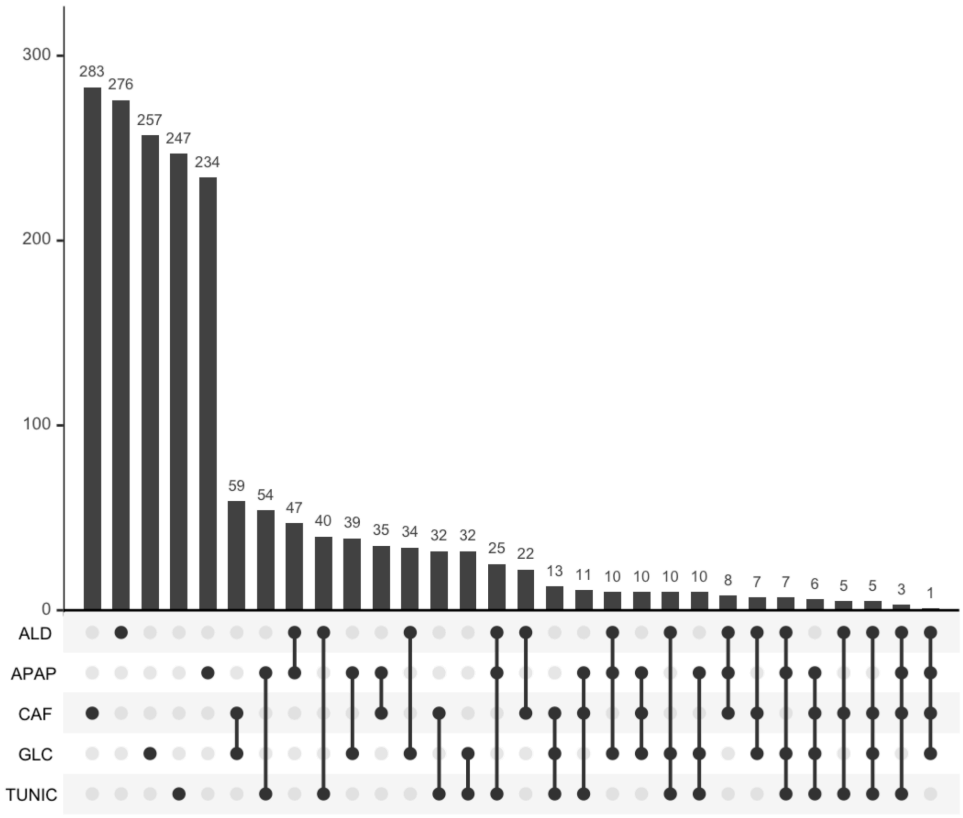


**Figure S7. Overlap between differentially expressed gene sets in LCLs.** Upset plot shows the overlap in gene identities for the top 500 differentially expressed genes in response to each treatment noted on the x-axis. Treatment acronyms as in Figure 1.

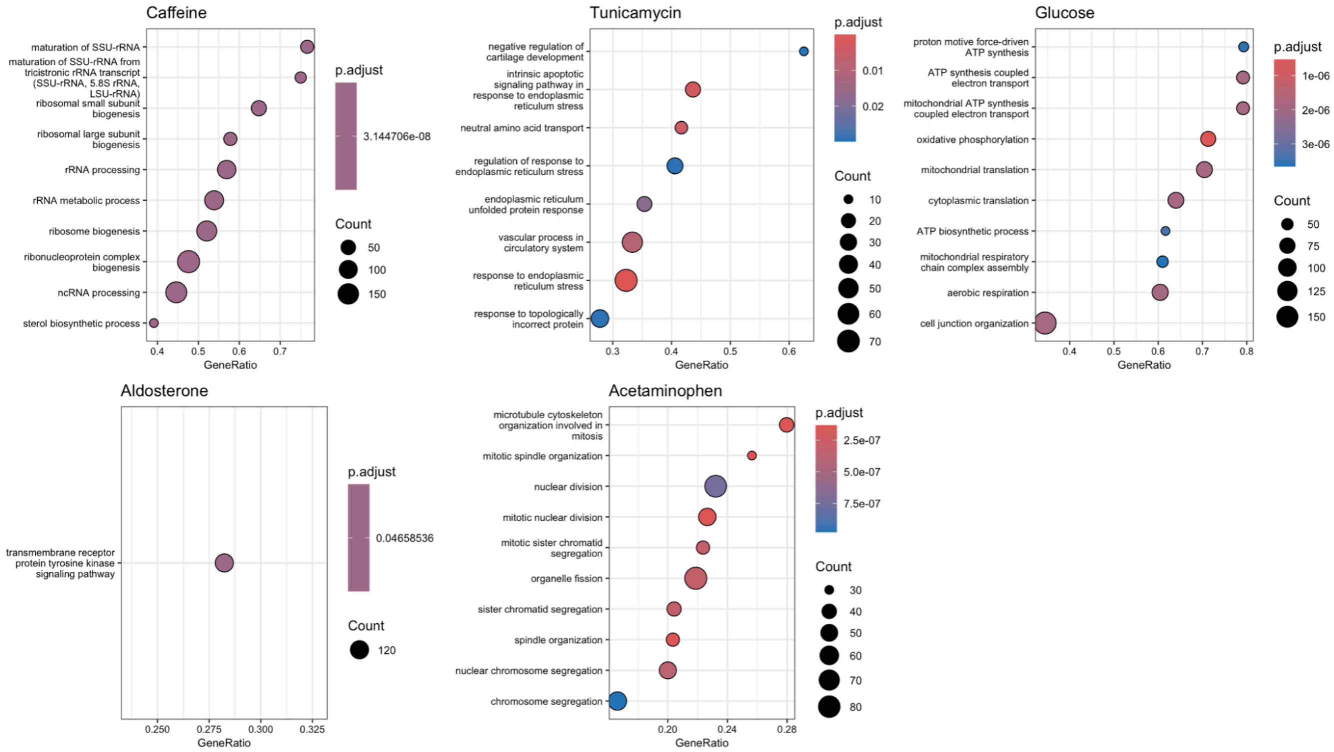


**Figure S8. Gene ontology analysis of perturbation-induced changes in LCL gene expression.** Each panel shows the top 10 biological pathways, ranked by enrichment score, for expression change following stimulation with the perturbation shown in the panel title(gene ontology enrichment analysis, all adjusted p-values<0.05). Gene Ratio was calculated by the clusterProfiler package in R and corresponds to the proportion of query genes that mapped to a specific biological pathway.

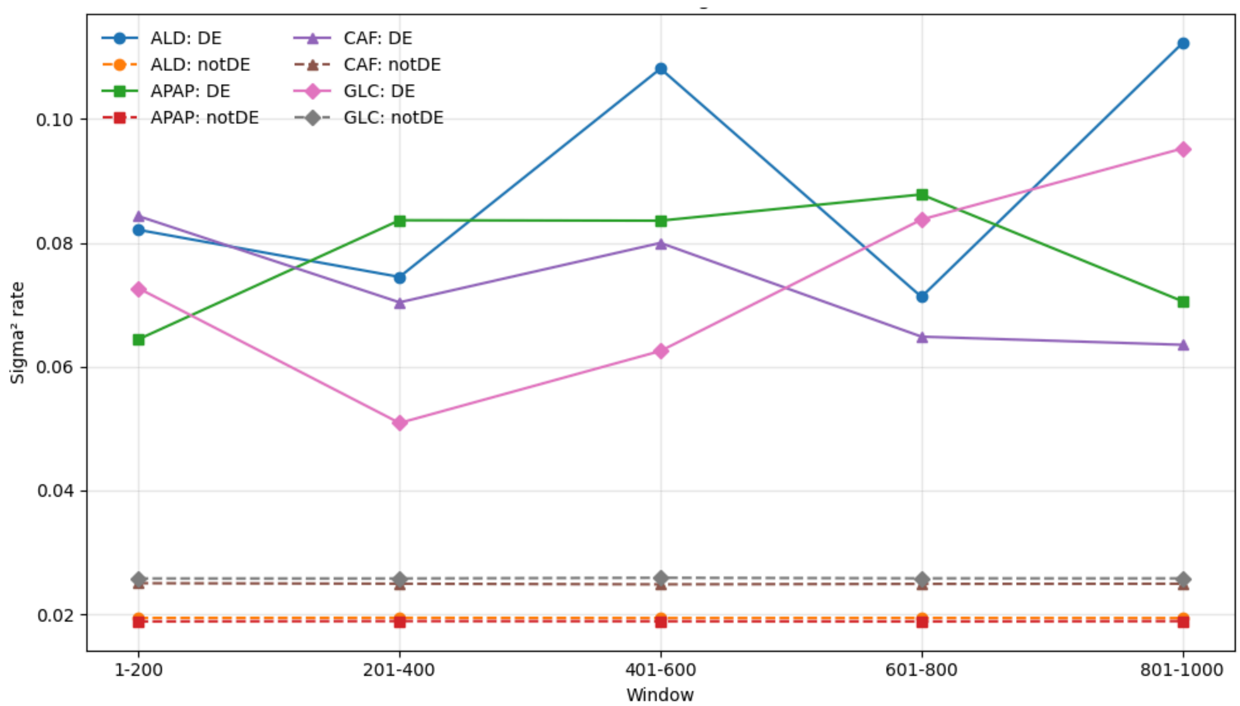


**Figure S9. Results from sliding window analyses**. We performed a sliding window analysis where we estimated and compared the rates of evolution for the top 1-200, 201-400, 401-600, 601-800, 801-1000 DE genes (rank-ordered by their effect sizes) to the estimated rate for the bottom 1000 DE genes. We observed similar evolutionary rate differences across the sliding windows.

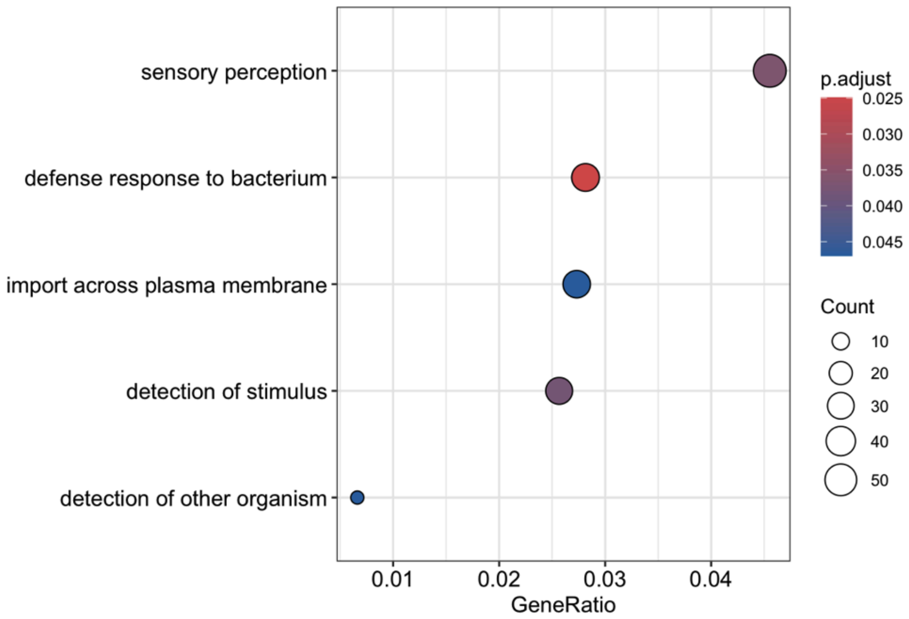


**Figure S10. Gene ontology analysis of significantly accelerated genes in white blood cells.** Plot shows all biological pathways that exhibited significant enrichment within genes with accelerated evolutionary rates (in any lineage) (gene ontology enrichment analysis, all adjusted p-values<0.05). Gene Ratio was calculated by the clusterProfiler package in R and corresponds to the proportion of query genes that mapped to a specific biological pathway. Gene expression data from [11].

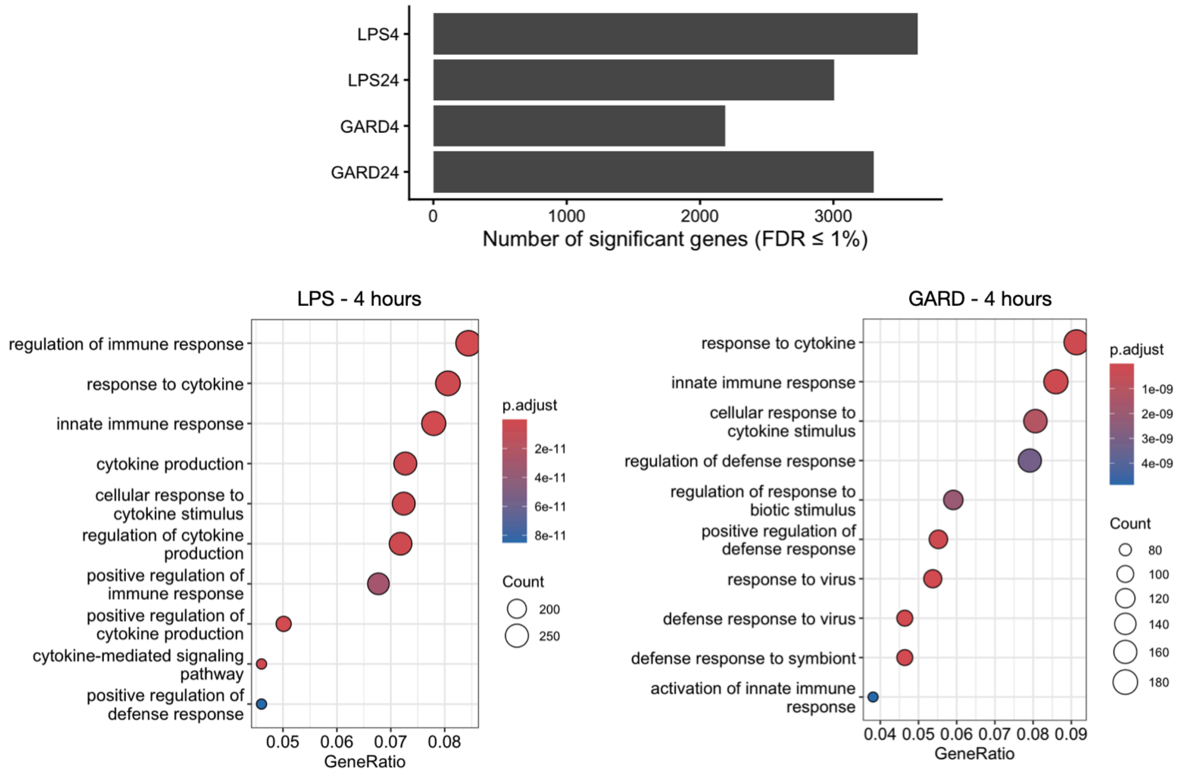


**Figure S11. Treatment responsive genes in white blood cells.** Barplot shows the number of differentially expressed genes in response to each treatment. Dotplots show the top 10 biological pathways that exhibited significant enrichment among differentially expressed genes (gene ontology enrichment analysis, all adjusted p-values<0.05). Gene Ratio was calculated by the clusterProfiler package in R and corresponds to the proportion of query genes that mapped to a specific biological pathway.

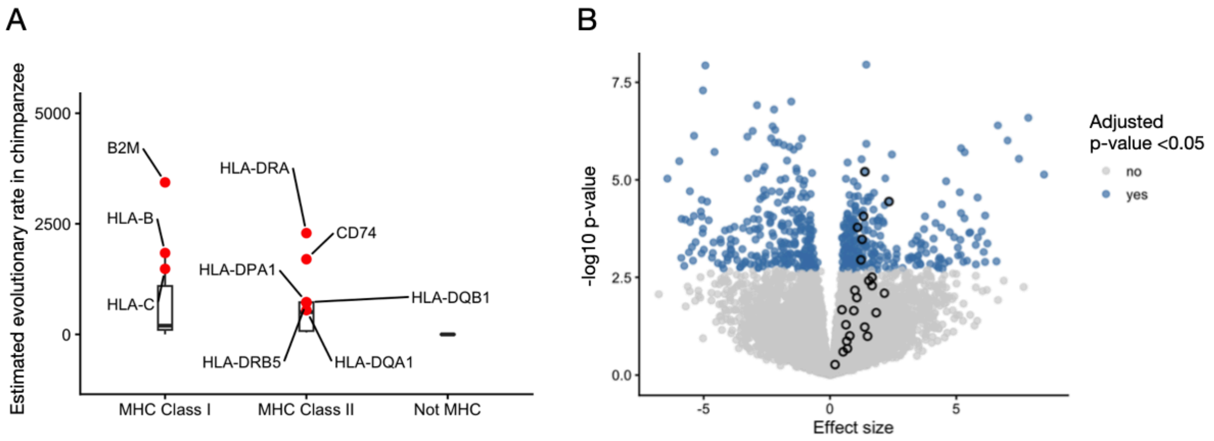


**Figure S12. Acceleration and differential expression of MHC genes in chimpanzees**. A) Boxplots of per gene estimates of evolutionary rates for genes included in the KEGG major histocompatibility complex (MHC) class I or class II molecule pathways (pathway IDs N00590 and N00363), versus all other genes. Red dots highlight significantly accelerated genes in MHC pathways with estimated rates >500. B) Volcano plot of results from a model comparing expression levels measured in the control condition between chimpanzees and all other taxa. Circles highlight MHC-related genes in the N00590 and N00363 KEGG pathways. All are upregulated (either significantly or non-significantly) in chimpanzees relative to other species.
